# Rapid molecular evolution of *Spiroplasma* symbionts of *Drosophila*

**DOI:** 10.1101/2020.06.23.165548

**Authors:** Michael Gerth, Humberto Martinez-Montoya, Paulino Ramirez, Florent Masson, Joanne S. Griffin, Rodolfo Aramayo, Stefanos Siozios, Bruno Lemaitre, Mariana Mateos, Gregory D.D. Hurst

## Abstract

*Spiroplasma* are a group of Mollicutes whose members include plant pathogens, insect pathogens, and endosymbionts of animals. *Spiroplasma* phenotypes have been repeatedly observed to be spontaneously lost in *Drosophila* cultures, and several studies have documented a high genomic turnover in *Spiroplasma* symbionts and plant pathogens. These observations suggest that *Spiroplasma* evolves quickly in comparison to other insect symbionts. Here, we systematically assess evolutionary rates and patterns of *Spiroplasma poulsonii*, a natural symbiont of *Drosophila*. We analysed genomic evolution of *s*Hy within flies, and *s*Mel within *in vitro* culture over several years. We observed that *S. poulsonii* substitution rates are among the highest reported for any bacteria, and around two orders of magnitude higher compared with other inherited arthropod endosymbionts. The absence of mismatch repair loci *mutS* and *mutL* is conserved across *Spiroplasma* and likely contributes to elevated substitution rates. Further, the closely related strains *s*Mel and *s*Hy (>99.5% sequence identity in shared loci) show extensive structural genomic differences, which potentially indicates a higher degree of host adaptation in *s*Hy, a protective symbiont of *Drosophila hydei*. Finally, comparison across diverse *Spiroplasma* lineages confirms previous reports of dynamic evolution of toxins, and identifies loci similar to the male-killing toxin Spaid in several *Spiroplasma* lineages and other endosymbionts. Overall, our results highlight the peculiar nature of *Spiroplasma* genome evolution, which may explain unusual features of its evolutionary ecology.

## Introduction

Many bacterial lineages have evolved to become associates of animal hosts (McFall-Ngai et al., 2013). In arthropods, such associations are the rule, and maternally inherited, endosymbiotic bacteria are especially common and diverse (Duron et al., 2008; Weinert et al., 2015). Prominent examples include obligate nutritional symbionts of blood and sap feeders (Douglas, 2009), reproductive manipulators (Drew et al., 2019), and protective symbionts (Oliver et al., 2014). Collectively, inherited symbionts of arthropods are highly diverse with respect to potential benefits and costs induced, and their degree of adaptation to hosts. For example, even within a single lineage of inherited symbionts - *Wolbachia* - there are nutritional mutualists (Hosokawa et al., 2010), protective symbionts (Teixeira et al., 2008), and reproductive manipulators (Dyson et al., 2002). Importantly, these symbiont-conferred traits may be modulated by environmental factors (Corbin et al., 2017) and host genetic background (Hornett et al., 2008).

*Spiroplasma* are small, helical bacteria that lack cell-walls. Like other Mollicutes (Mycoplasma-like organisms), they are exclusively found in host association and are often pathogenic (Razin, 2006). Among the first discovered spiroplasmas were plant pathogens (reviewed in Saglio and Whitcomb, 1979) and an inherited symbiont of *Drosophila* with the ability to shift sex ratios towards females (Poulson and Sakaguchi, 1961). Subsequently, *Spiroplasma* symbionts were found in a number of different *Drosophila* species (Haselkorn, 2010), as well as many other arthropods (Gasparich, 2002). More recently, molecular evidence was brought forward for *Spiroplasma* symbionts in non-arthropod animals (Duperron et al., 2013; He et al., 2018). In insects, two phenotypes induced by inherited *Spiroplasma* have been studied in detail (reviewed in Anbutsu and Fukatsu, 2011; Ballinger and Perlman, 2019). Firstly, *Spiroplasma* can enhance survival rates of its hosts under parasite or parasitoid attack. In *Drosophila neotestacea*, *Spiroplasma* inhibits growth and reproduction of the parasitic nematode *Howardula*, and thus reverses nematode-induced sterility (Jaenike et al., 2010). Further, *Spiroplasma* symbionts may greatly enhance *Drosophila* survival when attacked by parasitoids (Xie et al., 2010, 2014). Both protective phenotypes are mediated by *Spiroplasma* encoded RIP toxins (ribosome-inactivating proteins), which target the attackers’ ribosomes (Ballinger and Perlman, 2017; Hamilton et al., 2014, 2015). These processes may be complementary to protection through competition between *Spiroplasma* and parasitoids for lipids in the host hemolymph (Paredes et al., 2016). Secondly, *Spiroplasma* kills males early in development in several species of *Drosophila*, planthoppers (Sanada-Morimura et al., 2013), ladybird beetles (Tinsley and Majerus, 2006), lacewings (Hayashi et al., 2016), and butterflies (Jiggins et al., 2000). The male-killing phenotype is also linked to a toxin: in *Drosophila melanogaster*, a *Spiroplasma* protein containing OTU (Ovarian Tumour-like deubiquitinase) and ankyrin domains was demonstrated to kill male embryos, and thus termed *Spiroplasma poulsonii* androcidin (“Spaid”, Harumoto and Lemaitre, 2018).

*Spiroplasma* induced phenotypes were commonly observed to be dynamic compared with other symbiont traits, with repeated observation of phenotype change within laboratory culture over relatively short time frames. For example, spontaneous loss of male killing was found in *Spiroplasma* symbionts of *Drosophila nebulosa* (Yamada et al., 1982), and *Drosophila willistoni* (Ebbert, 1991), and spontaneous emergence of non male-killing *Spiroplasma* in *Drosophila melanogaster* has occurred at least twice in a single culture (Harumoto and Lemaitre, 2018; Masson et al., 2020). In addition, *Spiroplasma* symbionts artificially transferred from their native host *Drosophila hydei* into *D. melanogaster* initially caused pathogenesis, but evolved to become benign over just a few host generations (Nakayama et al., 2015). Genomic analysis has further documented dynamic *Spiroplasma* evolution driven by viral proliferation (Carle et al., 2010; Ku et al., 2013; Ye et al., 1996) and by extensive transfer of genetic material between plasmids, and from plasmids to chromosomes (Joshi et al., 2005; Mouches et al., 1984). Notably, plasmids in *Spiroplasma* commonly encode the systems that establish their phenotypic effect on their host (Berho et al., 2006). Examples include Spaid (Harumoto and Lemaitre, 2018) and RIP genes (Ballinger and Perlman, 2017), and rapid plasmid evolution may therefore contribute to the high phenotypic evolvability of *Spiroplasma*. Furthermore, elevated rates of evolution were found in various *Mycoplasma* species (Delaney et al., 2012; Woese et al., 1984), which are pathogens closely related to *Spiroplasma* (Gupta et al., 2019).

Bacteria adapting to hosts often follow similar evolutionary trajectories, and both increased mutational rates and proliferation of mobile genetic elements are commonly observed in diverse symbiotic taxa (McCutcheon and Moran, 2011). Therefore, spontaneous loss of *Spiroplasma* phenotypes and high genomic turnover could be explained by evolutionary mechanisms common to all symbiotic taxa. Although independent findings discussed above suggest that *Spiroplasma* symbionts may evolve quickly, this has never been quantified, and it has remained unclear how substitution rates compare to those of other symbionts. However, rates and patterns of evolutionary change may have important implications for *Spiroplasma* evolutionary ecology, and for its potential use in biological control programmes (Clark and Whitcomb, 1984; Schneider et al., 2019).

In this study, we systematically investigate rates and patterns of *Spiroplasma* evolution. To this end, we employed two strains of *Spiroplasma poulsonii*: 1) *s*Hy is a natural associate of *Drosophila hydei* (Ota et al., 1979) and confers protection against parasitic wasps (Xie et al., 2010). We monitored its evolution over ~10 years in fly culture, and determined changes through denovo reference genome sequencing and re-sequencing of multiple isolates. 2) *s*Mel occurs naturally in *Drosophila melanogaster*, where it kills male offspring (Montenegro et al., 2005) and protects from parasitoids (Xie et al., 2014). Using the previously established complete genome (Paredes et al., 2015) and cell-free culture (Masson et al., 2018), we re-sequenced isolates spanning ~2.5 years of evolution. This approach enabled tracing *Spiroplasma* evolution on various levels: within-strain (<10 years divergence), between strains, and through comparison with other *Spiroplasma* genomes, across a whole clade of arthropod symbionts. It further allowed comparison between symbionts evolving in a host and in axenic culture. We observed that rates of molecular evolution and chromosomal rearrangements in *Spiroplasma poulsonii* are substantially faster than in other inherited bacteria, and that this has resulted in markedly different genomic organisation in *s*Hy and *s*Mel. Our work thus highlights *Spiroplasma poulsonii* as an amenable, highly evolvable, but potentially unpredictable model insect symbiont.

## Materials and Methods

### *Spiroplasma* strains and genome sequencing

We documented the evolution of the *Spiroplasma poulsonii* strains *s*Hy and *s*Mel in fly hosts and in cell free culture, respectively (Tab. 1). sHy is one of the *Spiroplasma* symbionts naturally associated with *Drosophila hydei* (Mateos et al., 2006). Our evolution experiments covered ca. ten years and three variants of *s*Hy, all of which all originated from a single isolate (Fig. 1). Over the course of the study, *Drosophila hydei* hosts were maintained on standard corn meal agar diet (1% agarose, 8.5% sugar, 6% maize meal, 2% autolysed yeast, 0.25% nipagin), at 25°C, and a 12h:12h light:dark cycle. To isolate *Spiroplasma s*Hy-Liv DNA from *D. hydei*, we followed a protocol by Paredes et al. (2015): We collected ca. 300 *D. hydei* virgins and aged them for 4 weeks. The flies were then anesthetized with CO_2_ and pricked into the anterior sternopleurum using a sterile needle. Hemolymph was extracted via centrifugation (10,000g, 5 minutes, at 4°C) through a 0.45µm filter (Corning Costar Spin-X). To avoid hemolymph coagulation, we only extracted batches of 20 flies at a time and immediately transferred extracted hemolymph into ice cold phosphate buffered saline solution (1X PBS buffer; 137 mmol NaCl, 2.7 mmol KCl, 10 mmol Na2HPO4, 1.8 mmol KH2PO4). Extracts of all batches were combined, and the PBS solution with *Spiroplasma* cells was centrifuged (12,000g, 5 minutes, at 4°C). Supernatant PBS was discarded and the cell pellet was subjected to DNA extraction using a phenol-chloroform protocol and subsequent ethanol precipitation. DNA was eluted in water, and sequenced at the Centre for Genomics Research at University of Liverpool. Libraries were prepared using the Nextera XT kit and sequenced as 2×250bp run on an Illumina MiSeq sequencer.

**Tab. 1:** *Spiroplasma poulsonii* strains investigated in this study. Note that we follow the naming scheme suggested by Ballinger & Perlman (2017).

| Name used in the current study |  | Natural host | Establishment in fly or cell-free culture | DNA extracted from, date | Also known as |
| --- | --- | --- | --- | --- | --- |
| **sHy** | sHy-Tx12 | *Drosophila hydei* | 01/2004 (Mateos et al., 2006) | Hemolymph extract, 03/2012 | Hap_1, TEN104-106 |
|  | sHy-Liv18a | *Drosophila hydei* | 06/2008 (copy of sHy-Tx12) | Hemolymph extract, 04/2018 | - |
|  | sHy-Liv18b | *Drosophila hydei* | 03/2014 (copy of sHy-Liv18A) | Hemolymph extract, 04/2018 | - |
| **sMel** | sMel-Ug16 | *Drosophila melanogaster* | 10/2016 (Masson et al., 2018) | Cell free culture, 4 time points 2016–18 | MSRO-Uganda |
|  | sMel-Br18 | *Drosophila melanogaster* | 07/1997 (Montenegro et al., 2000) | Whole fly, 03/2018 | MSRO-RED42 |

**Fig. 1:**
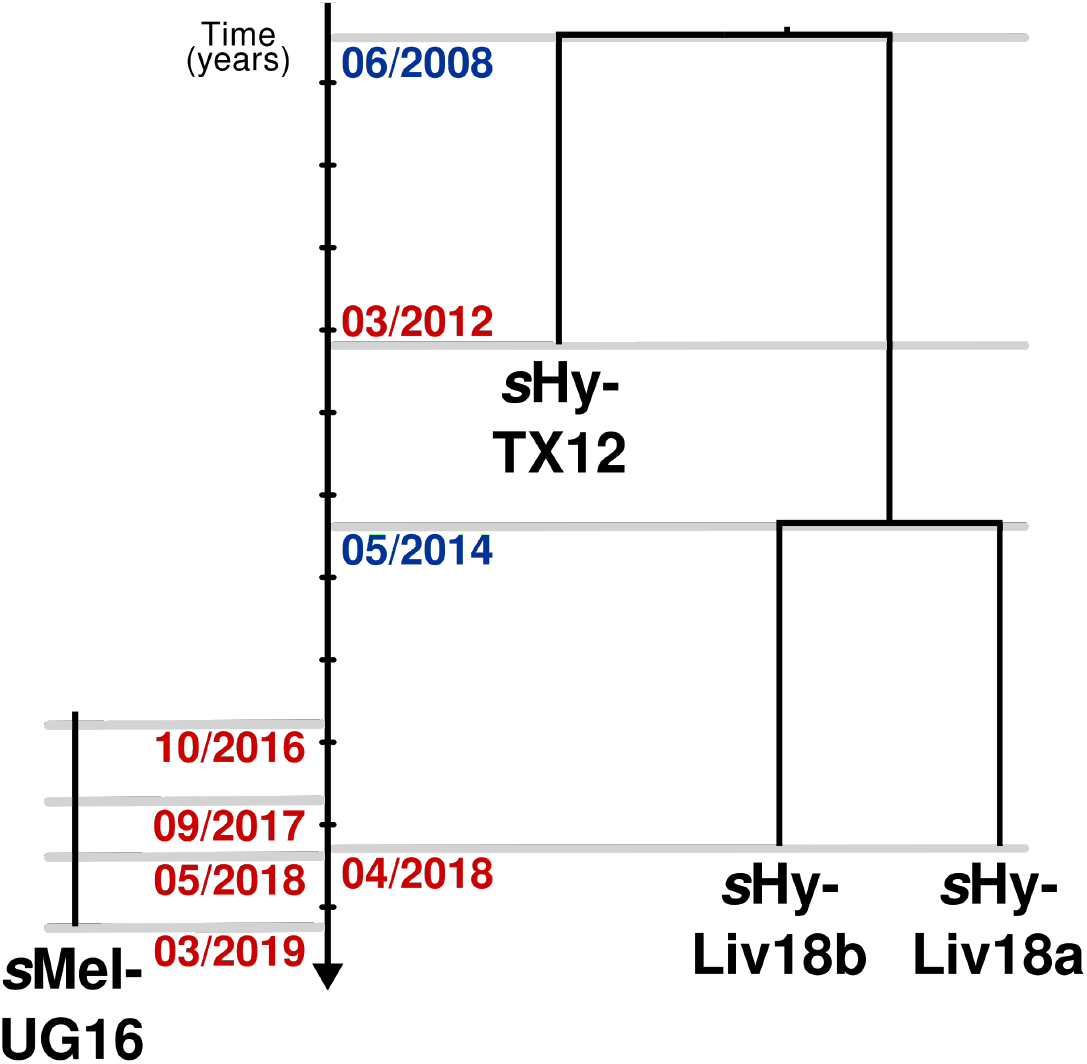
Overview on *Spiroplasma poulsonii* strains investigated in this study. Three *s*Hy strains were sampled from *Drosophila hydei* cultures over a total timespan of ten years. Blue dates denote splitting of *D. hydei* cultures, red dates mark *Spiroplasma* extraction for sequencing. For *s*Mel, four points of a time series of axenic culture were sequenced (HL - start of culture, P14 - after 14 passages, P57 - after 57 passages, P79 - after 79 passages). Here, red dates correspond to the date of isolation for sequencing.

DNA extraction for the *Spiroplasma* strain *s*Hy-Tx followed the protocol described above, with the following modifications: Immediately after the piercing, 35-40 flies were placed into 0.5 ml microcentrifuge tubes previously pierced in the bottom, which was placed within a 1.5 ml microcentrifuge tube containing 20µl PBS, and centrifuged at 4,500g for 10 seconds to collect hemolymph. DNA was recovered using a chloroform-ethanol procedure (Supplementary protocol 1) and diluted in AE buffer (Qiagen). A pair-ended library was constructed by Eureka Genomics (Hercules, CA). DNA was fragmented, end repaired, A’ tagged, ligated to adaptors, size-selected, and enriched with 25 cycles of PCR during which an index was incorporated to the sample. Sample preparation was performed according to Illumina’s Multiplexing Sample Preparation Guide and Eureka Genomics’ proprietary method. The resulting library was subjected to Illumina HiSeq 2500 sequencing at the Texas AgriLife Genomics and Bioinformatics Services Facility (College Station, TX).

*s*Mel is a naturally occurring *Spiroplasma* symbiont of *D. melanogaster* best known for its male-killing phenotype (Harumoto and Lemaitre, 2018; Montenegro et al., 2000). Cell free culture of *s*Mel-Ug was recently accomplished (Masson et al., 2018), and we used a time series of this culture to investigate *s*Mel molecular rates of evolution outside of host tissues. The time series covered 29 months in total, and sequencing was performed at four time points (Fig. 1). Illumina DNA sequencing was performed at MicrobesNG (Birmingham, UK) using the “standard service”. Culturing, passaging, and DNA extraction was performed as described in Masson et al. (2018).

We further sequenced a single isolate of sMel-Br deriving from a strain established in 1997 (Tab. 1). For Illumina sequencing, 1-2 grams of whole body flies from *Spiroplasma*-positive *D. melanogaster* were collected for CTAB-Phenol based DNA extraction. Illumina libraries were prepared with Illumina Truseq DNA kit and sequenced on an Illumina NovaSeq-6000 S2 150 paired end flow cell by Texas AgriLife Genomics and Bioinformatics Services Facility (College Station, TX). For Oxford Nanopore MinIon sequencing, heads were first removed by immersing whole flies in liquid Nitrogen, and vigorous shaking on a metal sieve. The headless specimens were immediately subjected to phenol-chloroform extraction. Nanopore library preparation was performed using the Ligation Sequencing kit 1D (SQK-LSK108), following precautionary measures to ensure long read lengths. The library was then sequenced on MinION sequencing device (MIN-101B) with an SpotON Flow Cell Mark 1 (R9.5-FLO-MIN107.1). Base Calling was performed using Albacore version 2.0 (Oxford Nanopore, Oxford, UK) and the reads were processed with Porechop version 0.24 (Wick et al., 2017a) to remove adapter sequences. A draft assembly of *s*Mel-Br was created using Illumina and Nanopore reads and Unicycler version 0.4.7 (Wick et al., 2017b). The assembly was fragmented, but did contain 3 circular contigs with sequence similarity to other *Spiroplasma* contigs.

### *s*Hy Reference genome sequencing and annotation

In order to reconstruct a high quality reference genome for *s*Hy, we extracted DNA from 30 *Drosophila hydei* virgin females carrying the symbiont. To limit the impact of potentially heterogeneous *Spiroplasma* populations within the flies, we used F3 individuals from a single isofemale line for DNA extraction. The flies were anesthetized at −20°C, washed with 70% ethanol, and briefly dried. The flies were then homogenized in G2 lysis buffer from the Qiagen Genomic DNA Buffer Set using a 1ml dounce tissue grinder (DWK Life Science), and DNA was extracted from the homogenate using a Qiagen Genomic-Tip 20/G following the protocol modifications from Miller et al. (2018). A total of 4μg DNA was then used in the “1D gDNA long reads without BluePippin protocol” library preparation protocol with the 1D Ligation Sequencing Kit (SQK-LSK108, Oxford Nanopore, Oxford, UK). Sequencing was performed on a MinION sequencer (Oxford Nanopore, Oxford, UK) using a single FLO-MIN106 R9 flow cell for 48 hours, and the raw Nanopore signals were base-called using Albacore version 2.3.1.

All reads passing the quality checks were used to create an assembly with Unicycler version 0.4.7 (Wick et al., 2017b). The resulting assembly contained 6 circular contigs (~15kb – ~1.6Mbp) with high similarity to the previously published *s*Mel genome as determined by BLAST+ version 2.9 searches (Camacho et al., 2009) in Bandage (Wick et al., 2015). We also created a hybrid assembly with Unicycler using all long reads and the *s*Hy-Liv18a Illumina reads. The hybrid assembly was more fragmented, but did contain 3 contigs which spanned almost the entire 1.6Mbp circular contig of the long-read only assembly. Because hybrid assemblies are typically more accurate than assemblies based on long reads only (De Maio et al., 2019), we corrected the long read assembly using the hybrid contigs. This was achieved by calling differences between the assemblies with the ‘dnadiff’ function implemented in MUMmer version 3.18 (Kurtz et al., 2004), and correcting the assembly accordingly. Finally, Pilon version 1.23 (Walker et al., 2014) was used to polish the corrected assembly, again using the *s*Hy-Liv18a Illumina reads. After nine rounds of polishing with Pilon, no further improvements could be observed and the assembly was considered final.

### Rates of molecular evolution in *s*Hy and *s*Mel

We determined changes in *s*Hy and *s*Mel chromosomes over time by employing the Snippy pipeline version 4.1.0 (Seemann, 2015). We used our newly assembled *s*Hy genome and the previously sequenced *s*Mel genome (Masson et al., 2018) as references, annotated using the Prokka pipeline version 1.13 with standard parameters and the *Spiroplasma* genetic code (translation table 4, Seemann, 2014). *s*Hy and *s*Mel variants determined by Snippy were filtered to only include positions which were covered by all *s*Hy or all *s*Mel Illumina libraries with at least 5x coverage, respectively. Further, to compare rates of molecular evolution between *Spiroplasma* and other microbial species, we counted the number of changes in third codon positions of coding sequences (CDS). For both strains, we only considered CDS that were entirely covered by sequencing depth of at least 5x in all Illumina libraries. Rates of molecular evolution were calculated by: number of observed changes at 3rd codon positions / number of considered 3rd codon positions / time in years over which the changes were observed. For *s*Hy, using *s*Hy-Liv18a as reference, we calculated two rate estimates, one based on comparison with *s*Hy-Liv18b, and one by comparing *s*Hy-Tx12 to the reference (Fig. 1). For *s*Mel, the hemolymph extract from which the culture was established was used as the reference (‘HL’, Fig. 1), and three rate estimates were calculated based on the changes observed in the three sampled time points of the culture (Fig. 1).

### Comparative genomics

We first compared the newly assembled *s*Hy genome with the most closely related, fully sequenced *Spiroplasma* genome, *s*Mel. Both genomes were annotated using the Kyoto Encyclopedia of Genes and Genomes (KEGG) and BlastKOALA (Kanehisa et al., 2016, 2006). Orthologous protein sequences determined using OrthoFinder version 2.2.3 (Emms and Kelly, 2015). Synteny between the genomes was assessed through aligning the genomes with Minimap2 version 2.15 (Li, 2018). Genome degradation was investigated by documenting the number of unique KEGG numbers for *s*Hy and *s*Mel, all of which were verified by BLAST+ searches. For each strain, we considered genes to be likely pseudogenized or truncated if the longest CDS within an orthogroup spanned at most 60% of the CDS length from the other strain.

Secondly, we compared *s*Hy with 17 *Spiroplasma* genomes from the Citri-Chrysopicola-Mirum clade (Table S1, Supplementary material for full list of names and accession numbers). All genomes were annotated using Prokka version 1.4.0, and orthology between predicted CDS was established using OrthoFinder. We extracted single-copy orthologs present in all strains, and aligned each locus separately with the L-INS-i method implemented in Mafft version 7.450 (Katoh and Standley, 2013). Recombination was evaluated with pairwise homoplasy index and window sizes of 100, 50, and 20 amino acids for each locus (Bruen et al., 2006), and any locus showing evidence for recombination was excluded. We used IQ-TREE version 1.6.10 (Nguyen et al., 2015) to reconstruct a core genome phylogeny from the remaining 96 loci (covering 26,019 amino acid positions). Each locus was treated as a separate partition with a distinct evolutionary rate (Chernomor et al., 2016), and optimal models and number of partitions was estimated with IQ-TREE (Kalyaanamoorthy et al., 2017; Lanfear et al., 2012). Branch support was assessed using 1,000 replicates of ultrafast bootstraps (Hoang et al., 2018), and approximate likelihood ratio test (Guindon et al., 2010) in IQ-TREE.

For all genomes, insertion sequence elements were annotated with Prokka. Prophage regions were predicted with PhiSpy version 3.7.8, which uses typical prophage features ((e.g., gene length, strand directionality, AT/GC skews, insertion points) for its predictions (Akhter et al., 2012). We also used the Phaster web server, which uses sequence similarities and other criteria to predict prophages (Arndt et al., 2016). We screened for the presence of toxin genes implied in protective phenotypes (Ballinger and Perlman, 2017; Hamilton et al., 2015) and male killing (Harumoto and Lemaitre, 2018). To this end, we downloaded UniProt sequence alignments from the Pfam database (Finn et al., 2016) for the protein families RIP (PF00161, ribosome inactivating protein) and OTU (PF02328, OTU-like cysteine protease), respectively. We used these alignments as databases for searches with HMMER version 3.2.1 (Eddy, 2011), and protein sequences from all genomes as queries. Domain architecture of all matching proteins was then determined using PfamScan with an e-value of 0.001 (Madeira et al., 2019), SignalP 5.0 (Almagro Armenteros et al., 2019), and TMHMM 2.0 (Krogh et al., 2001). RIP domains showed a high degree of divergence, and were therefore aligned to the reference RIP HMM profile using hmmalign from the HMMER software package. From this alignment, positions present in fewer than three sequences were removed, and a phylogeny reconstructed with IQ-TREE. OTU domains were aligned using MAFFT, and a phylogeny reconstructed with IQ-TREE.

Thirdly, we determined plasmid synteny across *Spiroplasma poulsonii* strains. In addition to the plasmids of *s*Mel and *s*Hy, we also included a plasmid of *s*Neo and all circular contigs from our *s*Mel-Br assembly. Plasmids were annotated using Prokka version 1.13 with a protein database composed of all plasmid proteins available on NCBI GenBank. Plasmid alignment and visualisation was performed with AliTV version 1.06 (Ankenbrand et al., 2017).

For an extended set of 31 *Spiroplasma* genomes that also included strains from the Apis clade (Table S1, supplementary material), we performed gene tree-species tree reconciliations to investigate the evolutionary history of prophage loci. To this end, we annotated all genomes and determined orthologous groups of loci as described above. Orthogroups were blasted against a database containing all available *Spiroplasma* virus proteins on NCBI Protein (N = 192). Genes were classified as ‘prophage related’ if hits were at least 60% identical over 50% of the length of any viral protein. Reconciliation was performed with the prophage loci using GeneRax version 1.2.2, using maximum likelihood gene trees determined with IQ-TREE as starting trees and the LG+G model for all loci. CRISPR/Cas systems and CRISPR arrays were predicted using the tools CRISPRCasTyper version 1.2.1 (Russel et al., 2020) and CRISPRIdentify (Mitrofanov et al., 2020), respectively.

### Data visualisation

Figures were prepared in R version 3.6.2 (R Core Team, 2016) using the packages ‘ape’ (Paradis and Schliep, 2019), ‘cowplot’ (Wilke, 2019), ‘ggalluvial’ (Brunson, 2019), ‘ggplot2’ (Wickham, 2009), ‘ggtree’ (Yu et al., 2017), and ‘ggridges’ (Wilke, 2020). Phylogenetic trees and domain architecture of toxin loci were visualised with EvolView version 3.0 (Subramanian et al., 2019).

## Results

### Rates and patterns of evolutionary change in *Spiroplasma poulsonii*

The *Spiroplasma poulsonii s*Hy reference genome comprises a single circular chromosome (1,625,797bp) and five circular contigs (23,069bp – 15,710bp) with sequence similarities to plasmids from other *Spiroplasma*. The chromosome contains 1,584 predicted coding sequences (CDS), a single ribosomal RNA cluster, and 30 tRNAs. Overall, genome content and metabolic capacities are highly similar to the previously sequenced, very closely related *s*Mel genome (Paredes et al., 2015), and a detailed comparison follows further below.

To estimate rates of molecular evolution in *Spiroplasma poulsonii*, we measured chromosome-wide changes in coding sequences of *Spiroplasma* from fly hosts (*s*Hy) and axenic culture (*s*Mel) over time. Our estimates for *s*Hy and *s*Mel are overlapping and range from 6.4×10^−6^ – 1.8×10^−5^ changes per position per year. This rate exceeds the rate reported for other symbiotic bacteria such as *Wolbachia* and *Buchnera* by at least two orders of magnitude (Fig. 2). Our estimate overlaps with rates calculated from some fast evolving human pathogens (e.g., *Enterococcus faecium* and *Acinetobacter baumannii*), and with evolutionary rates observed in the poultry pathogen *Mycoplasma gallisepticum* (which is closely related to *Spiroplasma*). Indeed, *Spiroplasma* substitution rates fall at around the lower estimates for RNA viruses (Fig. 2).

**Fig 2.**
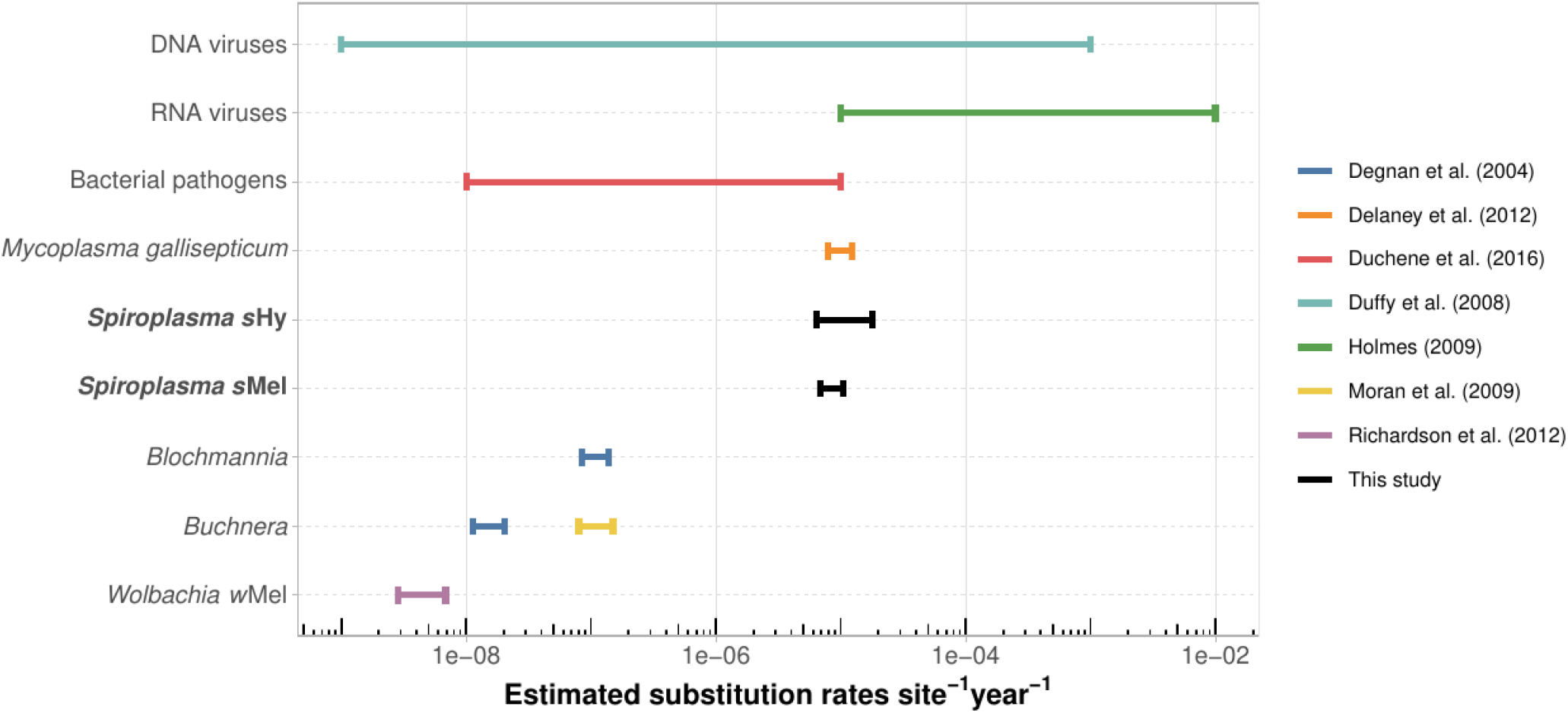
Comparison of estimated evolutionary rates across various microbes. Estimates obtained in this study are highlighted in bold. All bacterial rates are from chromosomal sequences only. DNA viruses - estimates as summarized in Duffy et al. (2008), including ssDNA viruses and dsDNA viruses. RNA viruses - range of RNA virus substitution rates from Holmes (2009). Bacterial pathogens - range of evolutionary rates estimated from genome wide data of 16 bacterial pathogens (*Acinetobacter baumannii*, *Bordetella pertussis*, *Enterococcus faecium*, *Klebsiella pneumoniae*, *Mycobacterium leprae*, *Mycobacterium tuberculosis*, *Neisseria meningitidis*, *Pseudomonas aeruginosa*, *Salmonella enterica*, *Shigella dysenteriae*, *Shigella sonnei*, *Staphylococcus aureus*, *Streptococcus pneumoniae*, *Streptococcus pyogenes*, *Vibrio cholerae*, *Yersinia pestis*) as determined by Duchêne et al. (2016). *Mycoplasma gallisepticum* - genome wide rate estimated from multiple isolates over a 12 year period (Delaney et al., 2012). *Blochmannia* and *Buchnera* (blue line) - based on 16S rDNA, *gidA*, and *groEL* sequences, taken from Degnan et al. (2004). *Buchnera* (yellow line) - genome-wide rate estimated from 7 isolates (Moran et al., 2009). *Wolbachia w*Mel - genome-wide data extracted from 179 *Drosophila melanogaster* SRA libraries (Richardson et al., 2012).

Over ~10 years of evolution, we observed a similar absolute number of variants on the chromosome (~1.6 Mbp) and plasmids (~0.1 Mbp) of *s*Hy, i.e., relatively more changes on the plasmids (Fig. 3, Table S3, supplementary material). Most variants in CDS affected hypothetical proteins, but were enriched in prophage-associated loci and adhesin related proteins (ARP, Fig. 3). Overall, *s*Hy variants in CDS were about equally often found to be synonymous as non-synonymous (Fig. 3) and changes were biased towards GC > AT substitutions (Fig. S1, Supplementary material). The changes in *s*Mel over ~2.5 years in culture affected only 15 different CDS in total, of which four were ARPs, and three lipoproteins (Table S2, supplementary material). Thus, the rates and patterns of evolutionary change are similar between the axenically cultured *s*Mel and the host associated *s*Hy.

**Fig. 3:**
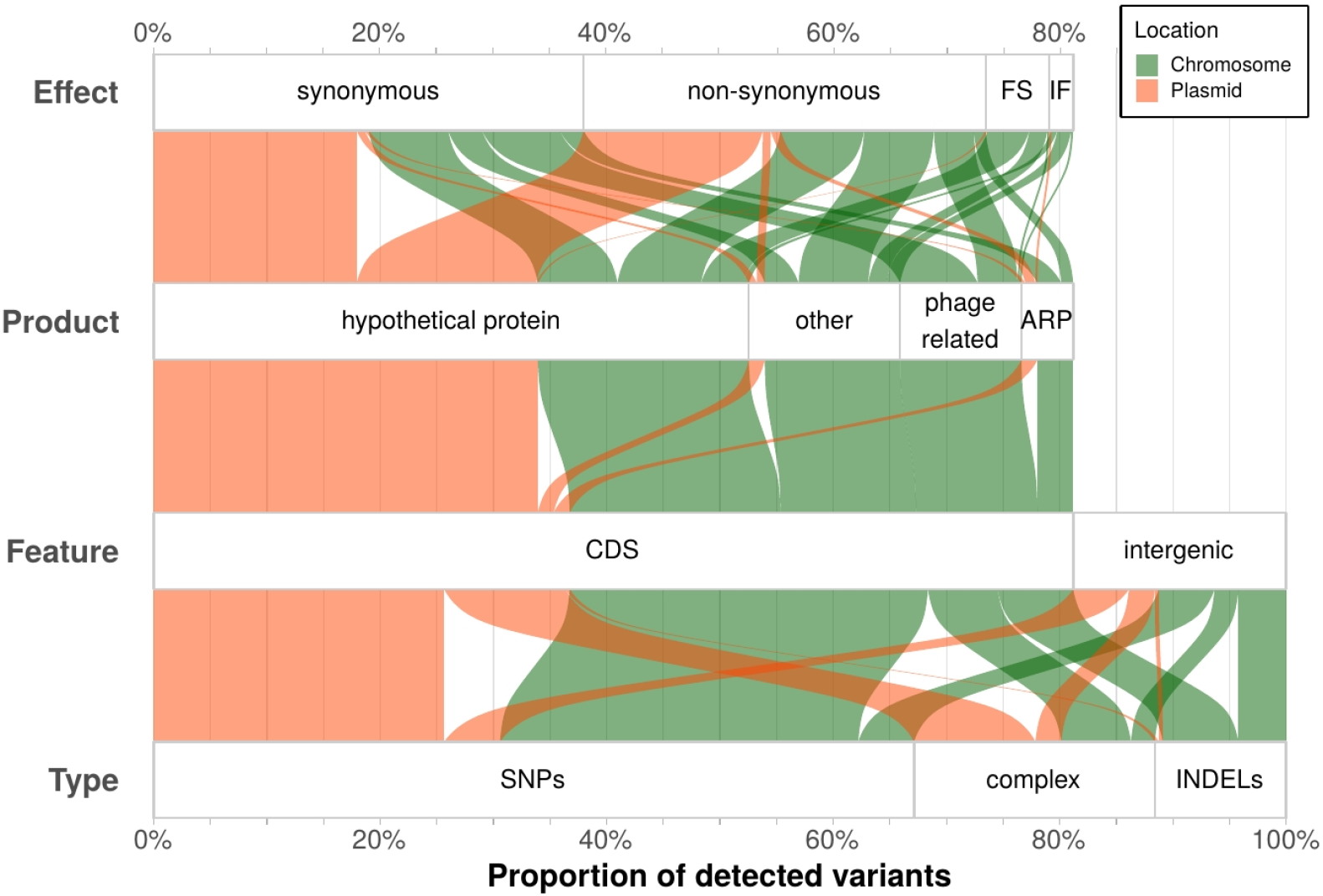
Overview of all observed variants in *s*Hy (100% = 570 variants). All annotations (Type, Feature, Product, Effect) are taken from results of the snippy pipeline. Abbreviations: ARP - Adhesion related protein, CDS - predicted coding sequence, complex – variants that span multiple nucleotides and/or SNPs, FS - frame shift, IF - in frame, INDELs - insertions and deletions SNPs - single nucleotide polymorphisms.

Comparing the genomes of *s*Hy and *s*Mel revealed a notable contrast between the high degree of nucleotide sequence identity on the one hand, and striking structural and gene content differences on the other hand. In 604 single copy orthologues shared between the genomes, average nucleotide identity was >99.5%. However, compared with *s*Mel, *s*Hy is reduced in gene content (1,584 vs 2,388 genes), chromosome size (1.61 vs 1.88 Mbp), and coding density (65% vs 82%). Further, *s*Hy contains fewer predicted insertion sequences (9 vs 89 in *s*Mel) and intact prophages as predicted with Phaster (*s*Mel: 16 such regions spanning 426 Kb; *s*Hy: 2 regions, 14 Kb), but not PhiSpy (Fig. S2, Supplementary material). Both *s*Mel and *s*Hy have a number of missing or truncated (i.e., potentially pseudogenized) genes when compared with each other, but the level of genomic deterioration is higher in *s*Hy, and covers a range of different genes (Fig. 4a, Table S4, supplementary material). Notable pseudogenised or absent loci in *s*Hy include parts of the phosphotransferase system, in particular the loci required for the uptake of N-Acetylmuramic Acid and N-acetylglucosamine (see Table S3, Supplementary material for a full list). Further, while recA is truncated in *s*Mel, the copy in *s*Hy appears complete and functional. As suggested by Paredes et al. (2015), the loss of recA function in *s*Mel is therefore likely very recent.

**Fig 4:**
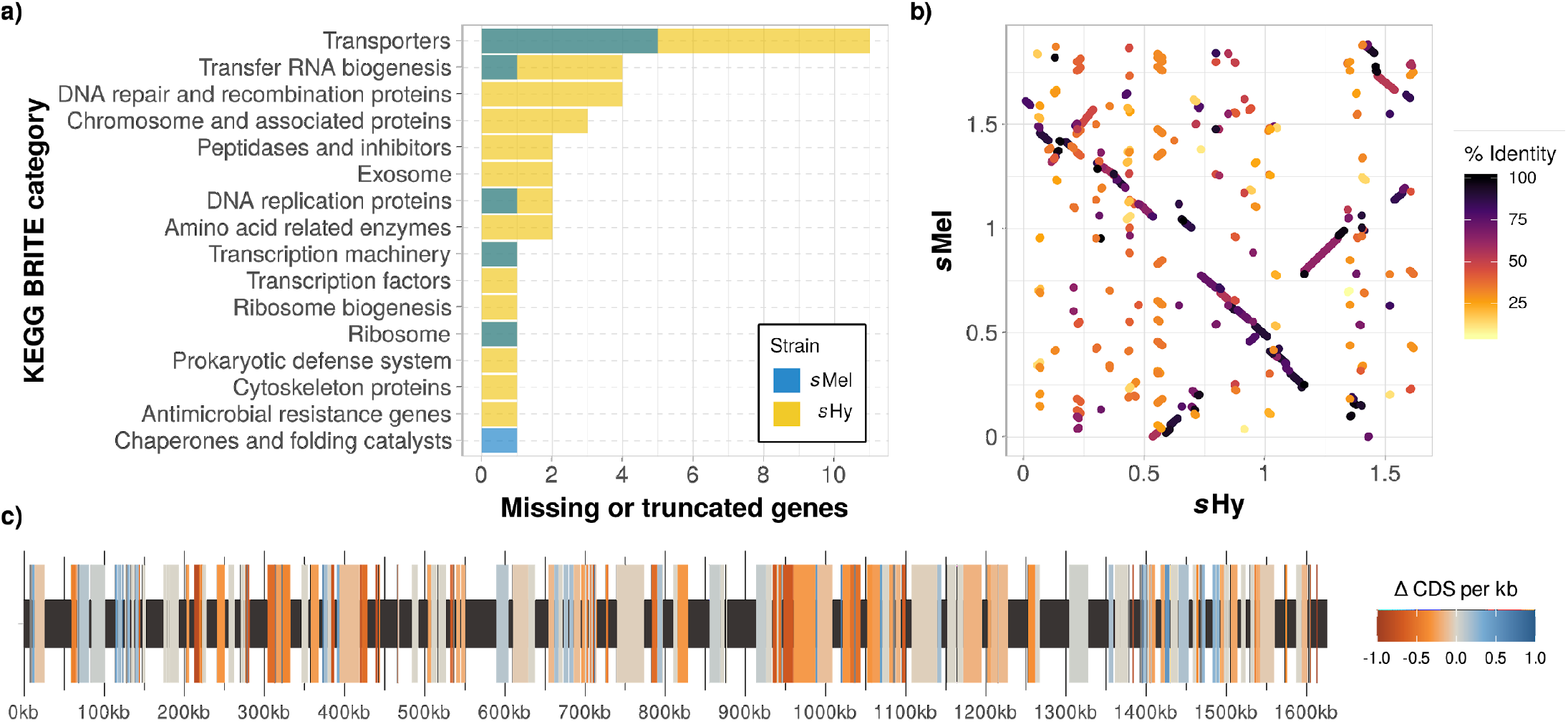
Comparison of genomic features between *s*Hy and *s*Mel. **a)** Number of genes with KEGG annotation that are present in one strain, but absent or truncated in the other. See Table S4, supplementary material for a complete list. **b)** Synteny as determined with minimap2. Syntenic between blocks are coloured by sequence similarity. **c)** Genomic map of *Spiroplasma poulsonii s*Hy with blocks syntenic to *s*Mel highlighted. The average number of predicted CDS per kb in these blocks is displayed as difference between *s*Hy and *s*Mel: positive and negative values indicate fewer CDS in *s*Hy or *s*Mel, respectively. Syntenic blocks were determined with a BLAST+ search using the *s*Mel chromosome sequence as query against a *s*Hy chromosome database, keeping a single best match for any *s*Hy region, and discarding hits below 1,000bp and under 95% nucleotide sequence similarity. Out of 635 syntenic blocks, 324 (spanning 852,730bp) contained fewer CDS in *s*Hy, and 164 (345,546bp) contained more CDS in *s*Hy, with 147 regions (334,084bp) showing no difference in CDS number.

Strikingly, there is evidence for a history of extensive chromosomal rearrangements since the last common ancestor of *s*Hy and *s*Mel, and genome-wide synteny between the strains is low (Fig. 4b). On average, syntenic blocks between the strains contained fewer predicted CDS for *s*Hy (on average −0.3 CDS/kb, Fig. 4c), in line with a higher degree of genomic deterioration in *s*Hy compared with *s*Mel. A comparison of plasmids across different *Spiroplasma poulsonii* strains (*s*Hy, *s*Mel-Ug, *s*Mel-Br, and *s*Neo) revealed similar gene content across the plasmids of different strains, but large differences in arrangement and number (Fig. S3, Supplementary material).

### Genome and toxin evolution in the genus *Spiroplasma*

Comparing *s*Hy with other sequenced strains of the *Spiroplasma* clades Citri, Poulsonii, Chrysopicola, and Mirum showed dynamic genome evolution within this genus of symbionts (Fig. 5). All investigated spiroplasmas have reduced genomes (~1.1–1.9Mbp), and the Poulsonii strains *s*Hy, *s*Mel and *s*Neo are among the strains with the largest main chromosomes (Fig. 5). Plasmid numbers range from 0–5, and *s*Hy has the highest number of plasmids of any of the investigated strains, although plasmid number is unclear for some of the genomes in draft status, and *S. citri* may have seven plasmids (Saillard et al., 2008). Prophage regions of varying sizes were predicted in all of the genomes (despite the lack of clear homologues to viral sequences in some of these, Ku et al., 2013). Reconciliation of prophage gene trees with the *Spiroplasma* species tree revealed that prophage proliferation has likely happened relatively recently, and repeatedly in the Citri and Poulsonii clades (Fig. S4, Supplementary material). Prophage loci are entirely absent, or very low in number, for *Spiroplasma* strains that harbour CRISPR/Cas systems, or remnants thereof (Fig. S4, Supplementary material). Further, *Spiroplasma* genome size correlated with the number of insertion sequences (Fig. 5, Fig. S5, Supplementary material). The distribution of CDS sequence lengths varies across the investigated genomes (Fig. 5), which may be explained by differences in proportion of prophage regions, level of pseudogenization, and assembly quality. Overall, our comparison indicates that *s*Hy may be more degraded than its closest relatives *s*Neo and *s*Mel, with a smaller chromosome size, fewer predicted CDS and mobile genetic elements, and the lowest coding density of these three (Fig. 5).

**Fig 5:**
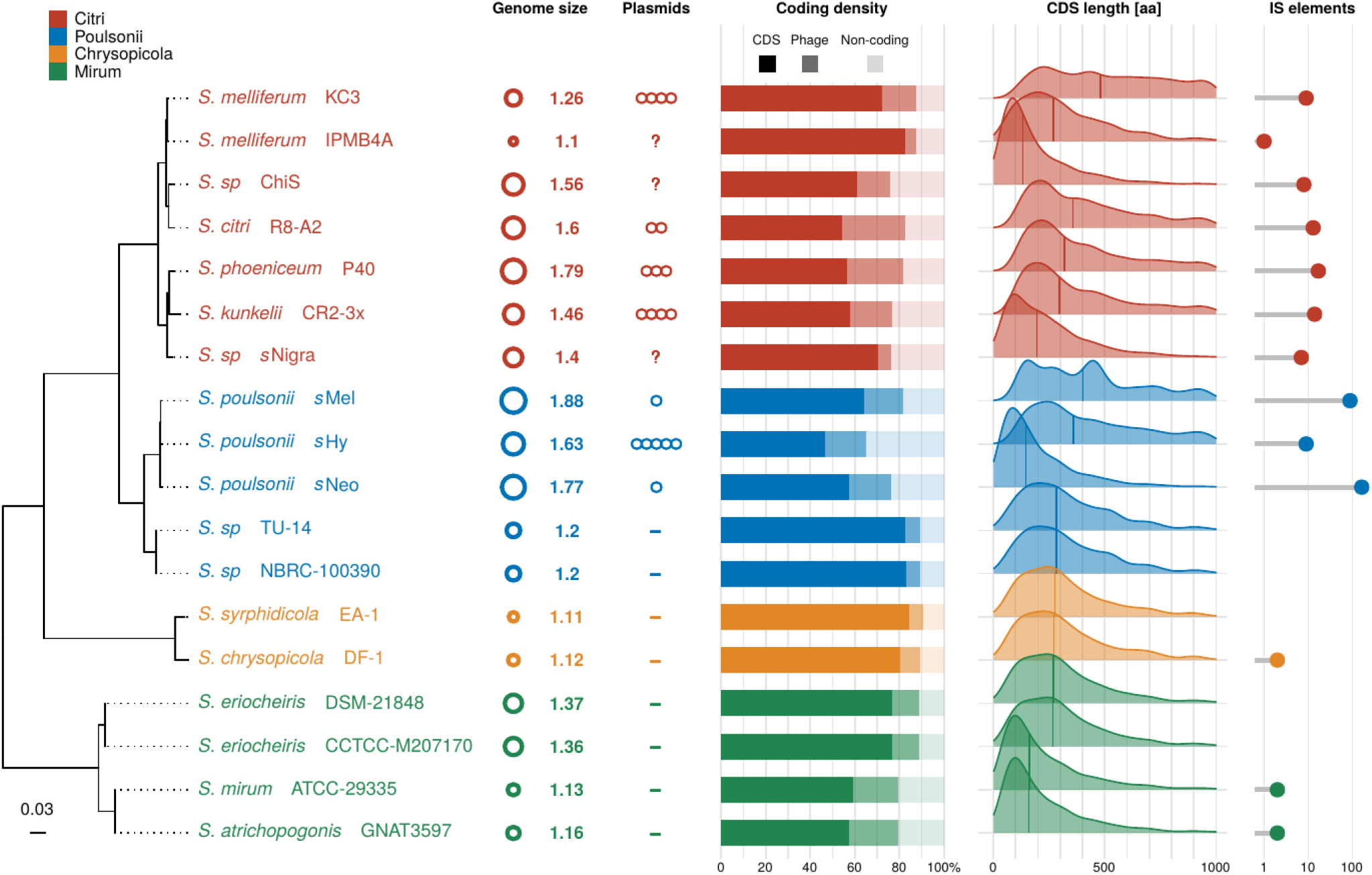
Genome properties of *Spiroplasma* strains from Citri, Poulsonii, Chrysopicola, and Mirum clades. Genome size and plasmid number was taken from the literature (Table S1, supplementary material). Coding density, CDS length and number of IS elements was determined with Prokka, and proportion of prophage was estimated with PhiSpy. *Spiroplasma* phylogeny is based on partitioned maximum likelihood analysis of 96 concatenated single copy genes present in all of the genomes (26,019 amino acid positions). All nodes were maximally supported by UFB and aLRT.

Spiroplasmas’ most striking phenotypes in insects have been mechanistically linked to toxin genes. For example, ribosome inactivating proteins (RIP) may protect *Spiroplasma* hosts by cleaving ribosomal RNA of parasites and parasitoids (Hamilton et al., 2015). Five RIP loci are present in *s*Mel (RIP1–5, of which 3–5 are almost identical copies). As expected, the protective *s*Hy also encodes RIPs, but only has a single orthologue for RIP1. However, it contains an additional RIP gene in three copies that appears to be absent in *s*Mel, and one of these copies also contains ankyrin repeats (Fig. S6, Supplementary material).

Further, the male killing phenotype of *s*Mel was recently established to be caused by *Spiroplasma* androcidin (Spaid), and both ankyrin repeats and a deubiquitinase domain (OTU) of this gene are necessary to induce male killing (Harumoto and Lemaitre, 2018). HMMER searches using OTU domain profiles revealed a number of *Spiroplasma* loci similar to Spaid (Fig. 6) which however lack its characteristic domain composition. For example, *s*Hy encodes three loci similar to Spaid: one lacks a signal peptide, one has no ankyrin repeats, and another encodes an epsilon-toxin like domain in addition to the OTU domain (Fig. 6). Other bacterial loci with notable similarities to the *Spiroplasma* OTU could only be detected in the symbiotic taxa *Rickettsia* and *Wolbachia*. In the phylogenetic reconstruction of OTU domains rooted with the eukaryotic sequences (from the ciliate *Stentor coeruleus*), *Rickettsia* and *Wolbachia* loci are nested within the *Spiroplasma* sequences. This topology suggests lateral gene transfer of OTU domain containing proteins from *Spiroplasma* to the other intracellular taxa, although differences in genetic code between *Spiroplasma* and other bacteria likely limit the probability for transfer of functional protein coding genes.

**Fig. 6:**
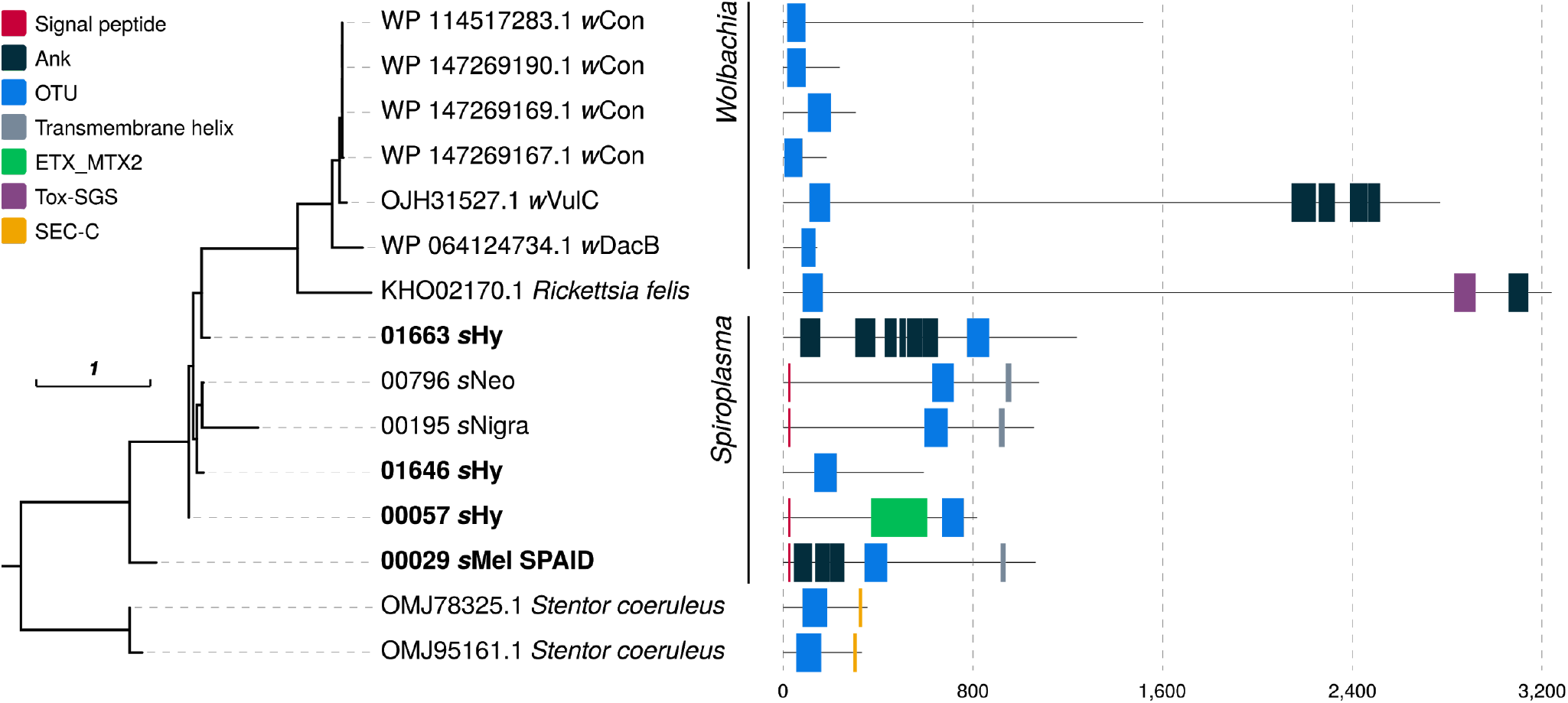
Spaid-like proteins in *Spiroplasma* and other symbionts. Maximum likelihood tree was reconstructed from an alignment of OTU domains (147 amino acid positions), and domain predictions are based on PfamScan, SignalP, and TMHMM. Abbreviations (corresponding to PFAM domains): Ank - ankyrin repeats, OTU - Ovarian tumor like deubiquitinase, ETX_MTX2 - Clostridium epsilon toxin ETX/Bacillus mosquitocidal toxin MTX2, Tox-SGS - Salivary gland secreted protein domain toxin, SEC-C - SEC-C nucleic acid binding domain.

## Discussion

### *Spiroplasma*, an exceptionally fast evolving symbiont

The substitution rates we observed in *Spiroplasma* symbionts are among the highest reported for any bacteria (Fig. 1). In a study comparing genome-wide evolutionary rates in bacterial human pathogens (Duchêne et al., 2016), most taxa showed considerably slower rates, and our estimate of *Spiroplasma* evolutionary rates overlaps with substitution rates of RNA viruses (Holmes, 2009). A number of reasons have been proposed for elevated substitution rates in bacterial symbionts and pathogens with small genome sizes: Firstly, host-associated microbes may acquire essential metabolic intermediates from their hosts and thus need not synthesise them (Moran, 2003). Obsolete metabolic genes subsequently accumulate now neutral substitutions, resulting in pseudogenization. Secondly, reduced population sizes of symbionts, together with bottlenecks at transmission events limit purifying selection of deleterious mutations (Moran, 1996) and mobile genetic elements (Lynch, 2006) – both in turn create further genomic deterioration. Thirdly, genomes of host-associated microbes are AT rich (Rocha and Danchin, 2002), which makes introduction of errors through replication slippage more likely (Moran et al., 2009). Fourthly, many symbionts lack DNA repair genes (Moran et al., 2008), which may result in increased substitution rates.

All of these points very likely apply for *Spiroplasma poulsonii*: its metabolic capacities are reduced compared with free living microbes, it has a high proportion of pseudogenes and prophages, very high AT content, and lacks several DNA repair genes. However, many symbiotic bacteria show similar trends of genome evolution (Burger and Lang, 2003), and it is therefore somewhat puzzling that *Spiroplasma* stands out with this exceptionally high substitution rate. Loss of DNA repair genes has been shown to co-occur with elevated mutation rates in intracellular bacteria (Itoh et al., 2002; Koonin et al., 1996; Massey, 2008), and may explain our observations. The loci encoding the mismatch repair proteins *mutS* and *mutL* are universally lacking in *Spiroplasma*, but are present in the slower evolving symbionts *Wolbachia w*Mel, and *Buchnera aphidicola* (Fig. 7, Richardson et al., 2012; Shigenobu et al., 2000; Wu et al., 2004). In *E. coli*, hypermutator strains that have lost such loci have a ~10– 300 fold increase of spontaneous mutation rates (LeClerc et al., 1996; Lee et al., 2012), which is comparable to the ~two orders of magnitude difference between substitution rates of *Buchnera* and *Spiroplasma* (Fig. 2). Like *Spiroplasma*, *Mycoplasma* universally lacks the mismatch repair loci *mutS*, *mutL*, and *mutH* (Carvalho et al., 2005; Chen et al., 2012). In line with this observation, elevated evolutionary rates were hypothesized for mycoplasmas (Woese et al., 1984), and in our comparison, *M. gallisepticum* appears to be the only bacterial taxon with substitution rates similar to *Spiroplasma* (Delaney et al., 2012). Absence of DNA mismatch repair pathway may thus be ancestral to Entomoplasmatales (Spiroplasmatacea + Entomoplasmataceae) and contribute to the dynamic genome evolution across this taxon (Lo et al., 2016; Rocha and Blanchard, 2002). Alternatively, increased substitutional rates caused by the loss of these loci could have arisen multiple times independently in Entomoplasmatales.

**Fig. 7:**
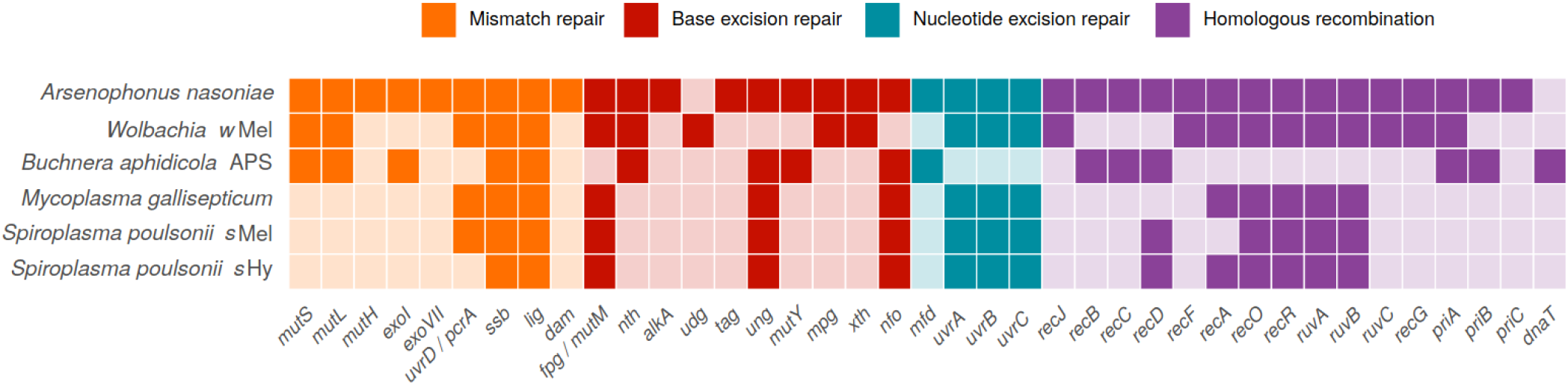
Absence (light colors) and presence (dark colors) of genes involved in DNA repair in *Buchnera aphidicola*, *Wolbachia w*Mel, *Mycoplasma gallisepticum*, and *Spiroplasma poulsonii s*Hy and *s*Mel. *Arsenophonus nasoniae* is included as an example of a symbiont with a relatively large genome and complete DNA repair pathways. Genes involved in more than one pathway are listed only once, and those missing in all of the taxa are not displayed. Data is taken from the KEGG pathway database (Kanehisa et al., 2006), and annotation of *s*Mel and *s*Hy genomes as described in Materials and Methods section.

Further to absence of DNA repair genes causing elevated mutation rates, a recent comparative study demonstrated a strong negative correlation between mutation rate and genome size in free living and endosymbiotic bacteria (Bourguignon et al., 2020). This correlation is however not apparent in the genomes of endosymbionts we have investigated. For example, the considerably slower evolving *Buchnera* genomes are much smaller than *Spiroplasma*, and *Wolbachia* would be predicted to have much larger genomes if their size was mainly determined by mutational rates. This suggests that substitution rates alone are a poor predictor for the sizes of here investigated genomes. Likely, these genome sizes result from an interplay of multiple factors such as population size, patterns of DNA repair gene absence, and mutational rates (Kuo et al., 2009; Marais et al., 2020).

Our rate estimate is potentially biased by at least two factors. Firstly, we have only investigated laboratory populations of *Spiroplasma poulsonii*. Each vertical transmission event creates symbiont population bottlenecks potentially increasing genetic drift and thus substitution rates (McCutcheon and Moran, 2011). Because the number of generations in natural populations of the *Spiroplasma* host *Drosophila hydei* is lower compared with laboratory reared hosts, vertical transmission events are rarer under natural conditions, and substitution rates therefore potentially lower. Further, laboratory strains could experience relaxed selection compared with natural symbiont populations. This may lead to higher substitution rate estimates from laboratory populations compared with natural populations. Secondly, substitution rates often appear larger when estimated over brief time periods (Ho et al., 2005). Duchene et al. (2016) found that substitution rates measured over 10 years can be up to one order of magnitude larger than those measured over 100 years. In agreement with such bias, we found more variants in our *s*Mel culture after 19 months than for the 29 months isolate (Table S3, supplementary material). The back mutations that likely have happened between 19 and 29 months of our *s*Mel culture would go unnoticed when comparing genomes over larger time scales.

More generally, it is difficult to estimate divergence times between *Spiroplasma* strains. The substitution rates estimated in *s*Hy and *s*Mel would suggest the two strains have diverged 1,120–2,260 years ago. However, this estimate is unreliable due to a number of factors that we cannot control for. For example, the number of generations per year for *Drosophila* hosts of *Spiroplasma* differ depending on species and location, and is expected to be lower in the wild compared with laboratory strains. Further, *Spiroplasma* may move between species, for example via ectoparasitic mites (Jaenike et al., 2007). A number of partially sympatric *Drosophila* species carry similar *Spiroplasma* strains (*D. hydei*, *D. melanogaster*, *D. willistoni*, *D. neotestacea*) (Haselkorn, 2010), and with our data it is impossible to determine the number and direction of potential *Spiroplasma* transfers between these species.

### Reductive genome evolution in *Spiroplasma*

According to a widely supported model of genome reduction in symbiotic bacteria (McCutcheon and Moran, 2011), the first stages of host restriction involve accumulation of pseudogenes and mobile genetic elements, chromosomal rearrangements, increased substitution rates, and excess deletions. Advanced stages of host association are accompanied by further genome shrinkage, and purging of mobile genetic elements, which overall result in more stable chromosomes. Using these characteristics, different levels of genome reduction are apparent in the investigated *Spiroplasma* genomes, across which genome sizes correlate positively with number of mobile genetic elements (plasmids, prophages, insertion sequences, Fig. 5, Fig. S5, supplementary material). On this spectrum, *S. syrphidicola* and *S. chrysopicola* have the most reduced genomes, without homologues of prophage loci, and high levels of synteny between the two strains (Ku et al., 2013). It was therefore argued that phages have likely invaded *Spiroplasma* only after the split of the Syrphidicola and Citri+Poulsonii clades (Ku et al., 2013). Our prophage gene tree-species tree reconciliations are in line with this hypothesis, but also indicate that prophage proliferation has largely happened independently in different *Spiroplasma* lineages (Fig. S4, supplementary material). CRISPR/Cas systems have multiple origins in *Spiroplasma* (Ipoutcha et al., 2019) and only occur in strains lacking prophages (Fig. S4, supplementary material). While the absence of antiviral systems often coincides with prophage proliferation (e.g., in the Citri clade), several strains with compact, streamlined genomes lack CRISPR/Cas and prophages (e.g., TU-14 & NBRC-100390, Fig. S4, supplementary material). These strains also show other hallmarks of reduced symbiont genomes (small size, high coding density, lack of plasmids and transposons, Fig. 5), which is in line with the model of genome reduction discussed above and suggests prophage regions were purged from these genomes. Alternatively, these strains may never have been exposed to phages.

Using the model of genome reduction during restriction to the host environment introduced above, *Spiroplasma poulsonii*’s genome characteristics suggest that such restriction has happened relatively recently. Although very closely related (99.5% sequence identity) and found in very similar hosts, *s*Mel and *s*Hy differ markedly in coding density, genome size, and proportion of prophage regions. Interestingly, PHASTER predicted 426 Kb of intact phage regions in *s*Mel, but only 14 Kb for *s*Hy (Fig. S2, supplementary material). Both phage proliferation in *s*Mel, and prophage loss in *s*Hy could have contributed to this. However, when using the similarity agnostic tool PhiSpy, the predicted prophage regions were similar in size between *s*Hy and *s*Mel (Fig. S2). This observation is compatible with degradation and/ or pseudogenization of prophage regions in *s*Hy, which would lead to reduced sequence similarity to viral loci (and thus reduced detectability by PHASTER), but not entirely blur prophage characteristics employed by PhiSpy (e.g., gene length, strand directionality, AT/GC skews, insertion points, Akhter et al., 2012). Since the split of the lineages, *s*Hy has not only lost prophage regions and insertion sequences compared with *s*Mel, but also several genes that are often lost in host restricted Bacteria, such as parts of the phosphotransferase system for the uptake of carbohydrates, one tRNA locus, and *uvrD* which plays a role in mismatch repair, and nucleotide excision repair (Table S2, supplementary material).

Using signatures of genomic degradation as a proxy, our findings collectively suggest that *s*Hy is in a more advanced stage of host restriction than *s*Mel. This may indicate co-adaptation with the host as a result of the fitness benefits associated with *s*Hy under parasitoid pressure, and the absence of detectable costs for carrying *s*Hy in *Drosophila hydei* (Osaka et al., 2013; Xie et al., 2010, 2014). However, the *Spiroplasma* symbiont of *Drosophila neotestacea s*Neo is also protective, does not cause obvious fitness costs (Jaenike et al., 2010), but has a less reduced genome (Fig.5, Ballinger and Perlman, 2017). Further, it is also possible that genome reduction in *s*Hy was mainly driven by stochastic effects or even by adaptation to laboratory conditions, as we have not investigated contemporary *s*Hy from wild *D. hydei* populations.

### Implications for *Spiroplasma* evolutionary ecology

*Spiroplasma* evolutionary ecology shows several parallels to that of the most widely distributed arthropod symbiont *Wolbachia*: both symbionts are found in a range of different hosts (Duron et al., 2008), have the ability to invade novel hosts (Conner et al., 2017; Haselkorn et al., 2009), may confer protection (Teixeira et al., 2008; Xie et al., 2010) but also kill males (Hurst and Jiggins, 2000), and can spread across host populations swiftly (Cockburn et al., 2013; Turelli and Hoffmann, 1991). However, there are also pronounced differences between the symbionts: *Spiroplasma* rarely reaches the high infection frequencies often observed in *Wolbachia* (Hilgenboecker et al., 2008; Watts et al., 2009), and is arguably found in a more diverse host range that encompasses arthropods and plants (Regassa and Gasparich, 2006), molluscs (Duperron et al., 2013), echinoderms (He et al., 2018), and cnidarians (Viver et al., 2017). Further, *Wolbachia* show greatly reduced spontaneous mutation rates compared with *Spiroplasma*, likely caused by a more complete set of DNA repair genes (Fig. 7).

In theory, fast evolutionary rates should enable *Spiroplasma* to adapt to novel hosts quickly (i.e., to reduce pathogenicity, and to maximise vertical transmission efficiency), and experimental studies have found high horizontal transmission efficiency of *Spiroplasma* (Haselkorn et al., 2013; Haselkorn and Jaenike, 2015; Nakayama et al., 2015). Consistent with this, we found that genes implicated in host-symbiont compatibility and virulence (e.g., adhesion-related proteins, have evolved especially fast in our evolution experiments. For example, adhesion-related proteins are important in cell invasion in other *Spiroplasma* species (Béven et al., 2012; Dubrana et al., 2016; Hou et al., 2017) and are enriched for evolutionary changes in *s*Hy and *s*Mel (Fig. 2). In addition, we documented dynamic evolution and turnover of toxin loci, which are important for host fitness and symbiont compatibility (Fig. 6, Fig. S6, supplementary material). This genomic flexibility may contribute to *Spiroplasma*’s broader host range when compared with *Wolbachia*. However, elevated evolutionary rates also make deleterious changes more likely, and, in the absence of strong selection, may result in faster loss of symbionts (Jaenike, 2012). This may explain the generally low prevalence of *Spiroplasma* symbionts (Haselkorn, 2010), which seems to increase only when carrying the symbiont is associated with a large fitness benefit (Jaenike et al., 2010). In contrast, virtually identical *Wolbachia* strains (“superspreaders”) are found in many different host species at very high frequencies (Turelli et al., 2018) – demonstrating that stationary genomes may be evolutionary advantageous. In summary, the nature of *Spiroplasma* genomic evolution likely contributes to its peculiar evolutionary ecology.

From a practical perspective, *Spiroplasma poulsonii* has many features of a desirable model for symbiont-host interactions: fast rates of evolution make it more likely that adaptation and spontaneous changes in phenotypes can be determined over short time scales, as has been observed previously (Harumoto and Lemaitre, 2018; Masson et al., 2020; Nakayama et al., 2015). On the other hand, fast evolutionary changes make experiments less predictable, and because stochastic effects become more pronounced, links of genomic changes with phenotypes may be obscured. Further, the generalisability of experimental results may be limited for extremely fast evolving symbionts. Our findings therefore also underline the importance of regular validation of laboratory symbiont strains through re-sequencing.

## Supporting information

Response to reviewers

## Author contributions

Designed and conceived project: MG, GDDH, MM. Experimental passage of *Spiroplasma* in vivo: MG, JG, GDDH, MM. Experimental passage of *Spiroplasma* in vitro: FM, BL. Symbiont DNA purification and sequencing: MG, PR, SS, RA, HMM, FM. Assembly and annotation: MG, HMM, PR, RA. Molecular evolution analyses: MG. Wrote first draft of paper: MG, MM, GDDH. All authors contributed in revising the manuscript and have read and approved the final manuscript.

## Acknowledgements

MG has received funding from the European Union’s Horizon 2020 Research and Innovation Program under Marie Sklodowska-Curie grant agreement 703379 and from EMBO through a long term fellowship (ALTF 48-2015) co-funded by the European Commission (LTFCOFUND2013, GA-2013-609409). Sequencing was supported by an award from the University of Liverpool Technology Directorate Voucher Scheme, and by a Texas A&M Genomics Seed Grant.

## Data accessibility

Raw reads for sHy-Tx can be found under NCBI BioProject Accession PRJNA274591 (Biosample SAMN08102256, SRA SRR6326389). Reads from sMel-Br are stored under PRJNA507275 (SAMN10488582). All other reads are available under NCBI BioProject PRJNA640980. Alignments and supplementary protocol are available from Zenodo under the DOI 10.5281/zenodo.3903208.

## Supplementary material

**Figure S1.** Mutational bias in *s*Hy evolution. Rates were calculated by counting the corresponding numbers of snps and dividing by the total number of AT or GC positions in the *s*Hy genome.

**Figure S2.** Predicted prophage regions in *s*Hy and *s*Mel by PHASTER (blue) and PhiSpy (yellow).

**Figure S3.** Plasmid synteny in *Spiroplasma poulsonii* strains. Numbers on ticks correspond to nucleotide position on plasmids in kb. Sequence similarities (>=60% and spanning >= 2kb) between plasmids are indicated by ribbons. Abbreviations: ARP - adhesion related protein, OTU - ovarian tumor domain containing protein (in *s*Mel: Spaid), ParA - plasmid partitioning protein like, RIP - ribosome inactivating domain containing protein, Spiralin - spiralin like protein. Figure was created with AliTV.

**Figure S4.** Prophage loci and CRISPR/Cas systems in *Spiroplasma*. **a)** Summary of prophage gene tree-species tree reconciliations. All events (gene speciations, duplications, transfers and losses) were inferred using GeneRax and are mapped onto the *Spiroplasma* phylogeny based on single copy orthologs present in all strains. **b)** CRISPR/Cas systems and arrays as predicted by CCtyper and CRISPRidentify, respectively. Maximum likelihood phylogeny is based on aligned Cas9 protein sequences, and was reconstructed using IQ-TREE. Note that CRISPR/Cas is absent in all Citri and Poulsonii strains. Several strains have lost or reduced parts of the CRISPR/Cas system, which therefore are likely not functional.

**Figure S5.** Correlations between genome size of investigated *Spiroplasma* strains and **a)** The number of IS elements as predicted by prokka; **b)**proportion of the genome encoding for prophages as predicted by PhiSpy. Abbreviations: R - Pearson’s correlation coefficients; p - p-values.

**Figure S6.** RIP loci in different *Spiroplasma* genomes. Maximum likelihood tree was calculated with IQ-TREE based on an alignment of RIP domains (288 amino acid positions), created using the hmmalign function of the HMMER software, and manually trimmed to exclude positions present in < 3 sequences. Domain prediction is based on PfamScan, SignalP, and TMHMM. Clades identified by Ballinger & Perlmann (2017) are indicated, and sequences from *s*Hy highlighted in bold font. UFB Bootstrap values ≥90 are shown on nodes. Abbreviations: SP - signal peptide, TM - transmembrane helix, Ank - ankyrin repeats, RIP - ribosome inactivating domain containing protein

**Table S1.** *Spiroplasma* genomes included in comparative genomic analyses, with accession numbers, genome properties, and phenotypes induced in host.

**Table S2.** Sequencing and mapping statistics for all *s*Hy and *s*Mel libraries used to infer variants.

**Table S3.** List of changes observed in *s*Hy and *s*Mel over the course of the evolution experiments.

**Table S4.** List of missing and truncated genes in *s*Hy and *s*Mel with KEGG annotation.

