## Supplementary figures and images for "Rapid molecular evolution of *Spiroplasma* symbionts of *Drosophila*"

### Figure S1

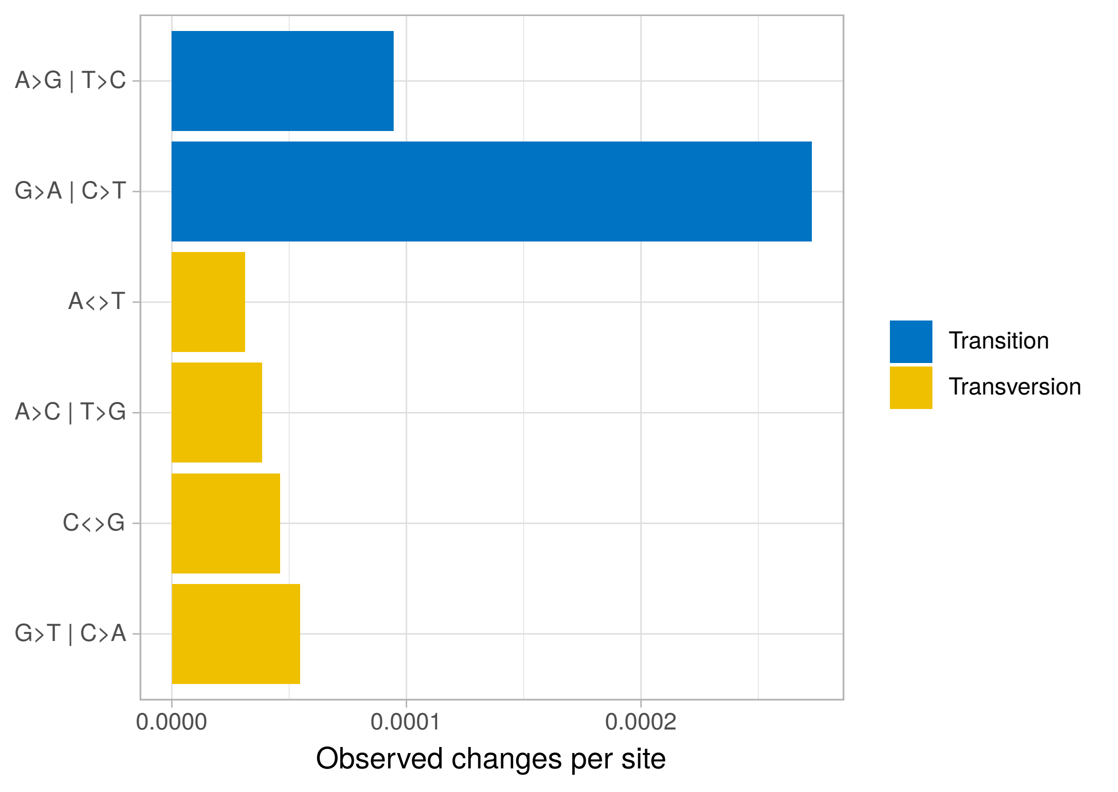

### Figure S2

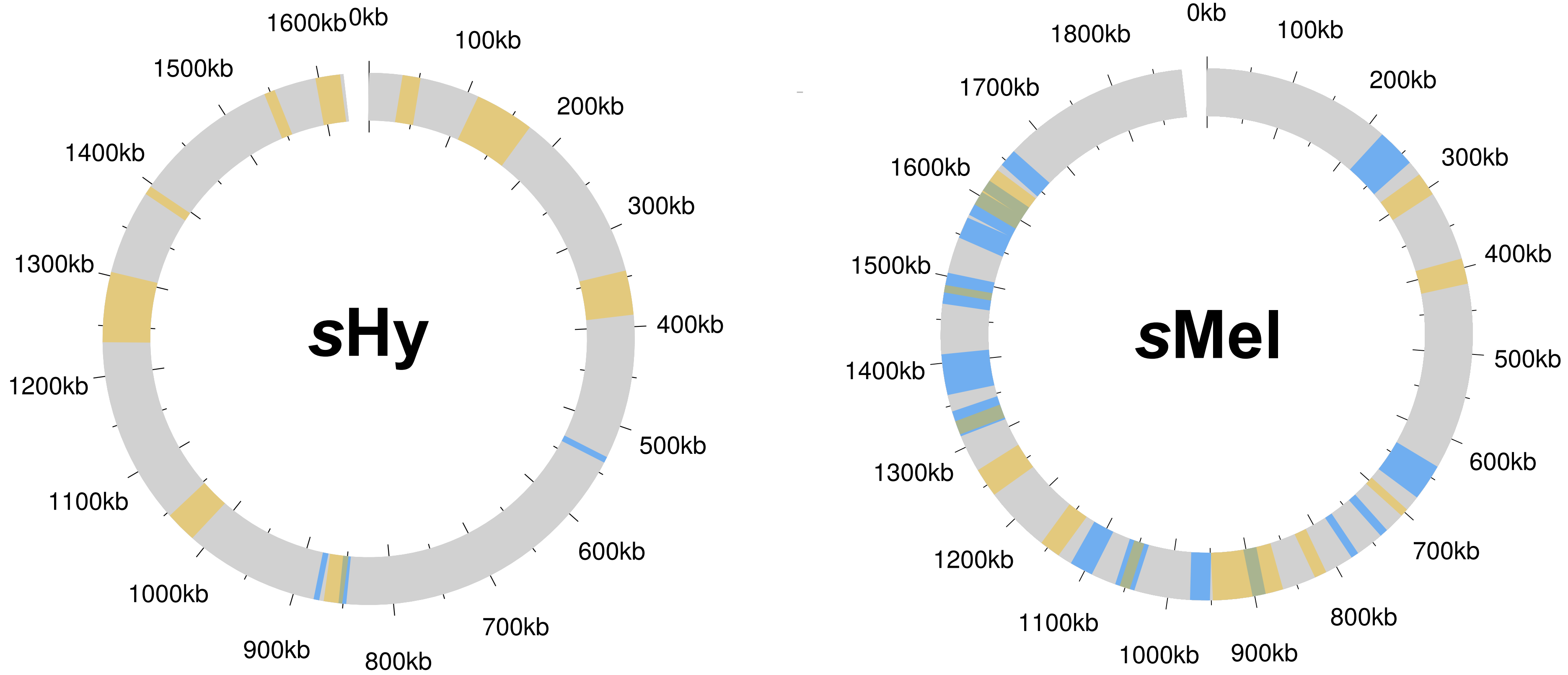

### Figure S3

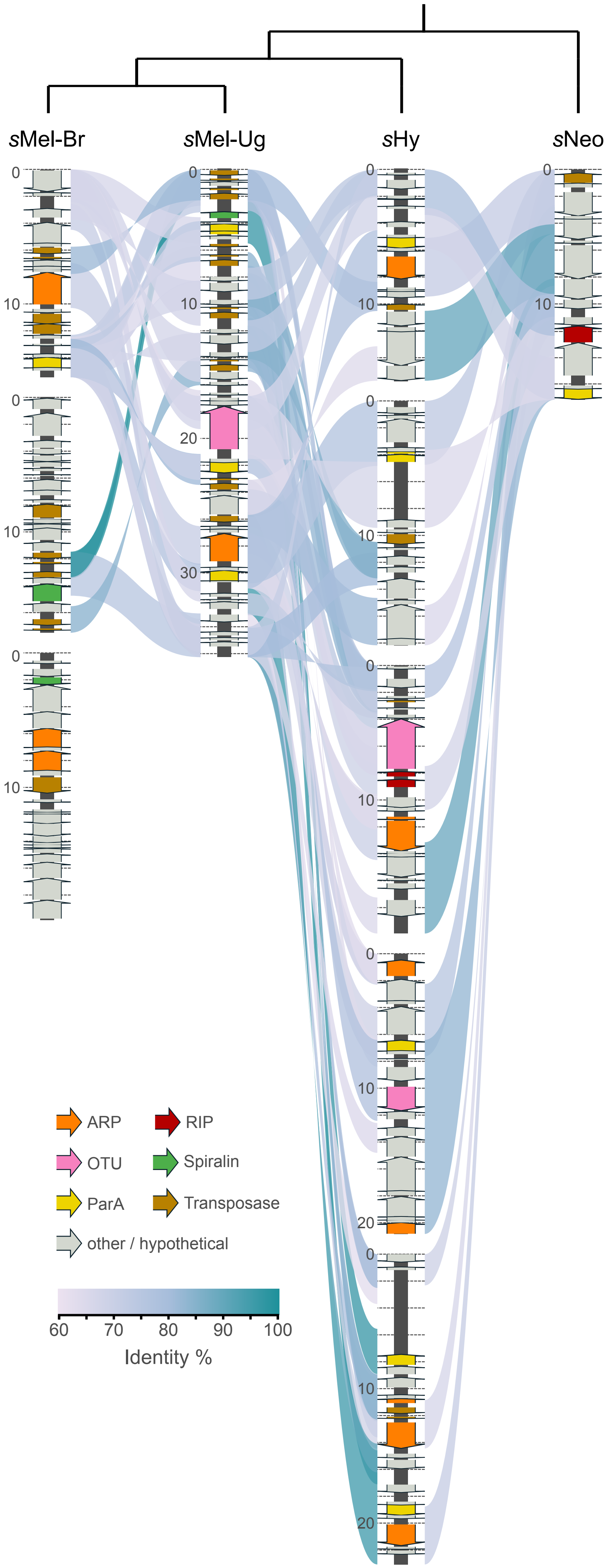

### Figure S4

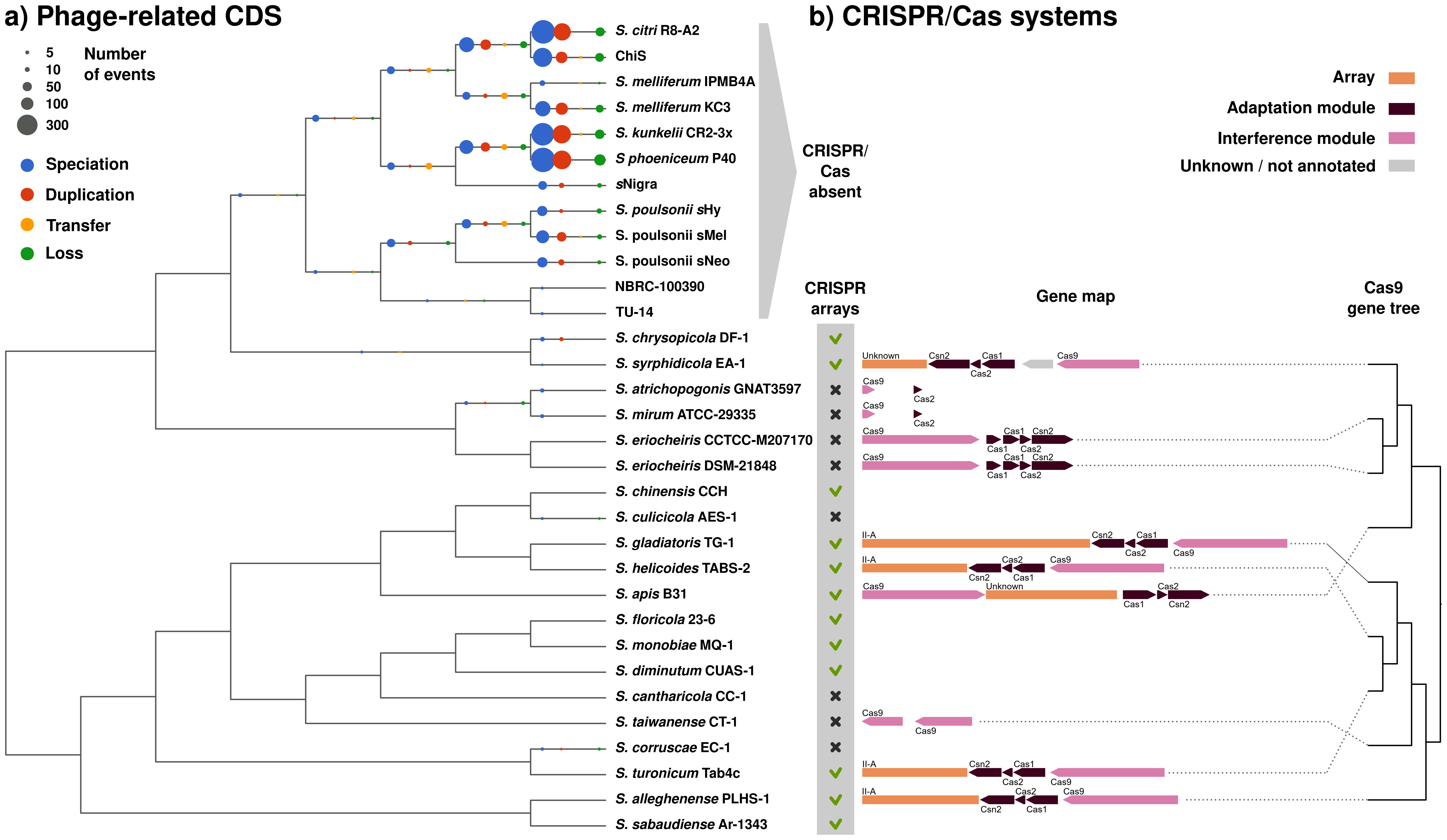

### Figure S5

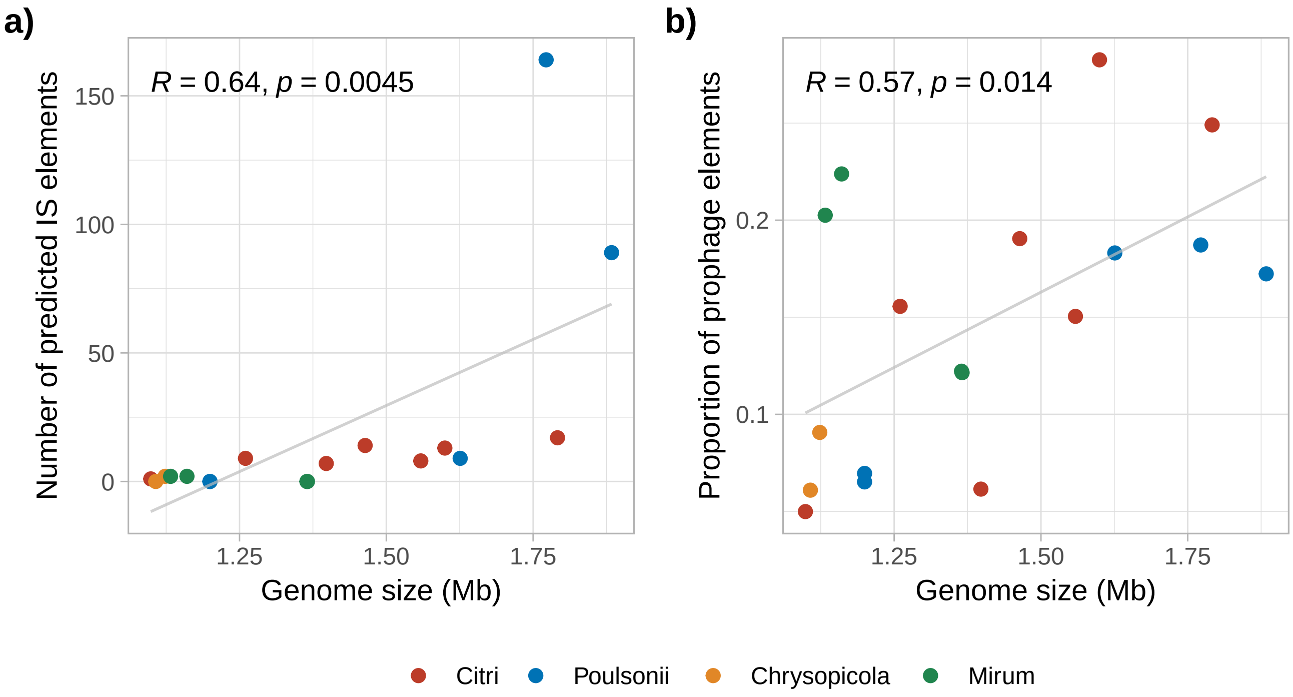

### Figure S6

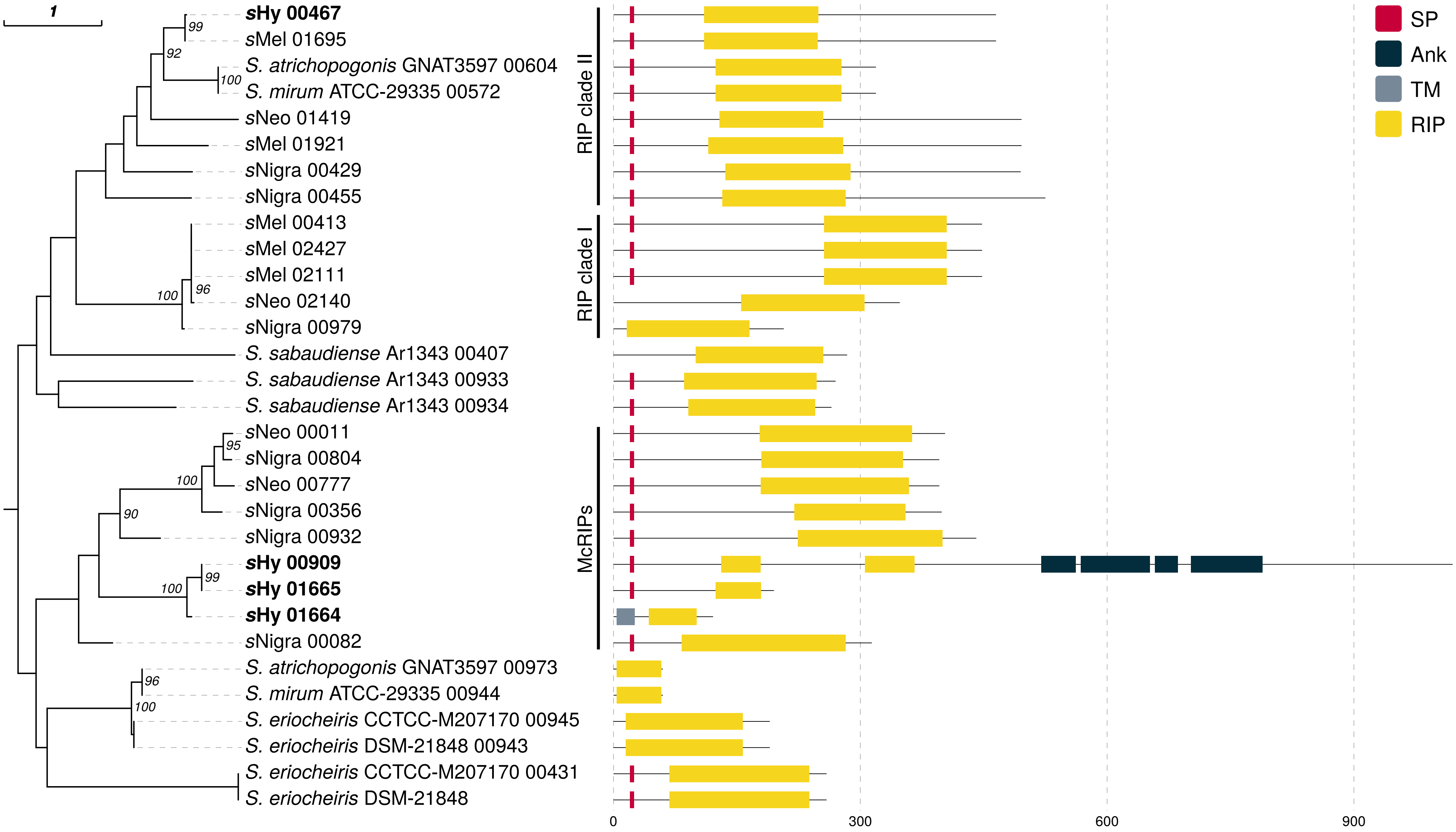
