## Supplementary material for "Rapid molecular evolution of *Spiroplasma* symbionts of *Drosophila*": Response to reviewers

### Reviewer #1:

The paper has potential. It's not there yet.

The paper presents a sequencing study describing the evolution of Spiroplasma over various years in lab cultures. Spiroplasma is a fascinating bacteria that induces some unique phenotypes including enhancing insect immunity or "protection" and male-killing. The premise for the study was that sometimes these phenotypes disappear in cultures and thus the bacteria is likely quickly evolving and subject to frequent mutation. The researchers sequence various cultures of Spiroplasma (sHy and sMel), assemble and annotate genomes, compare the genomes, quantify the rates of evolution and compare these rates to some other studies on viruses, human microbiota/pathogens, and wolbachia. They find that Spiroplasma evolve real fast and speculate that the mechanism for this is a lack of various Mut repair enzymes. They look at fast evolving proteins of interest including RIP toxins which kill nematodes and spaid which is an inducer of male killing. So essentially the big result here is that Spiroplasma evolves real fast.

In my opinion the paper is weak in a few senses. It doesn't reflect hypothesis driven science. It's mostly observational data and the researchers do not test any hypotheses. Now I don't think this is a deal breaker, but I do think it weakens the paper. Also, my comment should not imply that there isn't valuable data herein; and in fact I think the other big weakness is that the researchers do NOT exploit the true value of the data to derive and test novel hypotheses.

> We respectfully disagree with the reviewer's opinion that hypothesis driven papers are generally 'stronger' than observational studies. Arguably, valuable insights can be derived from both types of studies, and this has been discussed in depth elsewhere (e.g., https://doi.org/10.1186/s13059-020-02133-w). However, we did have a hypothesis when we designed this study, and it was based on previous reports that novel phenotypes occur commonly in *Spiroplasma* in lab culture. We hypothesised that molecular evolution of *Spiroplasma* is likely also very fast. We further conclude with novel hypotheses on the evolutionary ecology of *Spiroplasma poulsonii*.

For example: one aspect I was most excited about was to see how the researchers dissect and annotate evolutionary differences induced by axenic culture systems. The authors have the ability to compare and contrast genomes of Spiroplasma cultured in host insects AND Spiroplasma cultured without insects in axenic culture. Within these genome comparisons are likely novel insights that could shed light on mechanisms of maternal transmission and mechanisms of cell invasion etc… However, I was shocked to see that there is no in-depth analysis of specific proteins that are changing and evolving in these two diverse culture systems. I thought the analysis was entirely insufficient and didn't extract or present the real value of the datasets here. There are some brief mentions in the discussion of adherin binding proteins, but that was essentially it. I think the researchers focused too much on the past, ( the RIP toxins and spaid) rather than pointing out new interesting genes and hypotheses about them.

For example: Maternal transmission would no longer be required in axenic culture, what genes got mutated? This is perhaps the most interesting thing that is not even touched upon.

So essentially my main criticism is the added value from this paper which is the potential ability to compare symbiont genomes in hosts to symbionts with Axenic culture was NOT exploited. Given the novelty and impact of the axenic culture studies by Bruno, I would have hoped to see this upfront.

We agree in general that our dataset presents the opportunity to compare evolution of the symbiont in axenic culture and in the host. However, any potential interpretation of evolution in axenic culture vs. in-host is hampered by the fact that we were comparing two different strains of *Spiroplasma*. With a sample size of 1 each, any conclusions on evolution in axenic culture vs. in-host would have been speculative.

Additionally, we did not find notable differences in evolutionary rates or affected proteins between the two strains. From the first version of our paper:

"The changes in *s*Mel over ~2.5 years in culture affected only 15 different CDS in total, of which four were ARPs, and three lipoproteins"

– which is overall very similar to the changes observed in sHy (Fig. 3). We concluded that the same genes are likely to evolve in axenic culture and in the host. We have made this clearer now in the manuscript:

"The changes in *s*Mel over ~2.5 years in culture affected only 15 different CDS in total, of which four were ARPs, and three lipoproteins. [New version:] Thus, the rates and patterns of evolutionary change are similar between the axenically cultured *s*Mel and the host associated *s*Hy."

Also there are some paragraphs comparing broad genomic differences between sHy and sMel, but I didn't think the differences in how these genomes evolved over time in comparison to their earlier selves was emphasized or explained in enough detail.

We summarise the main patterns of change over time in *s*Mel and *s*Hy in the results and discussion sections, in Figure 3, and further list all detected changes from both strains in Supplementary table S2. We thus feel that the level of detail is appropriate here, especially given the length of the overall manuscript.

Another example of not exploiting the value of the data: The plasmids are usually where much of the action is in microbes. There should be detailed annotations and figures of the plasmids. Tell me what is on them. Tell me which genes are evolving. Tell me if there are operons. Tell me what pathways are in the plasmids. I found the discussions of plasmid results wholly lacking. I also inherently felt that discussions of plasmids should be kept completely separate from discussions of chromosome evolution, regardless of similar rates of evolution or not… Plasmids are unique selfish entities and I imagine their evolution is wholly distinct from the evolution of chromosomes. They deserve their own sections and figures (in my opinion).

> There is a figure comparing plasmid synteny and gene content across the investigated strains in the supplementary material. Notable loci are highlighted, and again, the majority of genes are uncharacterised.

The figure legends are completely insufficient and they ask me to read other papers to understand them, which is annoying.

> We apologise for this oversight and have now provided more comprehensive legends for all figures.

Other minor comments:

What about presence/absence of recA?

> recA is truncated in *s*Mel by a previous stop codon, as discussed in detail in Paredes et al. (https://doi.org/10.1128/mBio.02437-14). recA appears to be complete and potentially functional in *s*Hy, which supports Paredes et al's inference that the truncation in sMel may be relatively recent (prior to the split of sMel and sHy). The new version of the manuscript now includes this detail:
>
> "Further, while recA is truncated in sMel, the copy in sHy appears complete and functional. As suggested by Paredes et al. (2015), the loss of recA function in *s*Mel is therefore likely very recent."

There are differences in dna extraction prior to genome sequencing for each of the strains. I suspect this is because different individuals sequenced different genomes. But I worry that different protocols could produce different results and therefore a comparison might be tainted by dna extraction and library prep specifics. Can you at least explain to the reader why this is not an issue, if it is not an issue?

> DNA extraction procedures differed because they were done in different laboratories. All DNA extractions were based on phenol-chloroform, and all *Spiroplasma* extractions were based on isolating fly hemolymph. Any differences in protocols are minor, and mentioned mainly for reasons of reproducibility. We do not see any reason why this would affect genome reconstruction of a single bacterial isolate. Several studies suggest that the impact of DNA extraction and library preparation is negligible for assemblies and calling SNPs (e.g., https://doi.org/10.1016/j.heliyon.2019.e02745; https://doi.org/10.1038/s41598-020-71207-3).

Examples:

181 - why were heads removed? Why was this dna extraction protocol here different from the hemolymph extraction protocol? Might this have changed anything?

> Please see the comment on DNA extraction above. Head removal is often used when enrichment of symbiont DNA in whole fly extracts is desired.

195 - how much heterogeneity do you expect in any given fly. Do you have SNP data differences amongst good reads that could point out different alleles within a Spiroplasma population within an individual fly? It would be interesting to know which genes have a large amount of different alleles.

> As described in the methods section, we always pooled hemolymph from multiple fly individuals in order to extract sufficient DNA for genome sequencing, so we cannot say anything about the genetic heterogeneity of *Spiroplasma* populations in any single fly individual. The levels of heterozygosity in the pooled extracts were however very low: Out of all variants called with more than 10x coverage in sHy-Liv18B and sHy-TX12 strains, 98% and 95% were unanimously supported by all mapping reads, respectively. Only 0.8% and 1% of variants had 5% or more reads supporting an alternative allele, respectively. No alternative allele was supported by more than 18% and 11% of reads, respectively.

199 - another DNA extraction protocol. There isn't consistency here. If the reads and coverage are good enough, it shouldn't be a problem. But if there were data issues or assembly issues, this would raise concern in my mind. Can the researchers discuss or alleviate concerns here? Some assemblies have 6 chromosomes, some have 3 chromosomes. I presume these were different strains of Spiroplasma and not the same one?

> Please see the comment on DNA extraction above. As described in the methods section, we obtained long reads and short reads from the same DNA extract. Depending on the reads and algorithms employed, we created assemblies that differed in number of contigs. This is not unusual or unexpected (e.g., http://doi.org/10.1099/mgen.0.000132). A consensus was created by using a long read assembly and correcting it with contigs from a hybrid assembly, and subsequently, with Illumina reads. We feel that this was a good approach to ensure a contiguous, but accurate assembly.

Figure 1: were the samples that are 6 years apart (red) sequence in exactly the same way with the same technology? Could this produce any relics? Also, why display information for sMel in a table and information for sHy in a figure? Can't you creatively standardize a visual means of showing this information and compile information to one item?

Please see the comment on DNA extraction above. We have taken up the suggestion of the reviewer and created a single figure to display sampling for both strains.

I wonder what would happen if you took the same sample and did different DNA extraction protocols, different library prep protocols, and different illumina rounds of sequencing and independent algorithm assemblies… how much would they come out the same? Has anyone ever done this experiment? Is there any reference for this control that shows they would in fact come out the same? This is essentially what I am worried about here. This could be a minor issue, if the researchers could just confidently explain why this is NOT an issue.

Please see the comment on DNA extraction above.

Line 30 - you introduce sHy and sMel without defining what they are yet? Clarify immediately that they are both S.poulsoni

This was clearly stated in line 29 of our manuscript.

line 247 - They found fragmented genes with orthofinder, if it was less than 60% length homology… why set an arbitrary cutoff of 60? Anything less than 100 is possibly a pseudogenization if the last amino acid is important, or the C-terminus is important, which it often is… What is the rationale here?

We agree with the reviewer that this is a relatively crude measure of pseudogenization that likely results in missing several candidate pseudogenes. Because it is usually impossible to functionally characterise all loci of a bacterial genome, truncation is often used as an indication that genes may have lost their functions (https://doi.org/10.1093/nar/gki631). This limitation was acknowledged in the first version of the manuscript: “Both *s*Mel and *s*Hy have a number of missing or truncated (i.e., potentially pseudogenized) genes when compared with each other”.

To quantify an evolutionary rate, I read that they counted the number of changes in 3rd codon wobble positions/year. Why just wobble codons… why not all SNPs period? But then in the figure 2, it seemed like they are tallying a percentage of a total 100% = 570 "variants" or changes in the sequences (I wouldn't use the word variants, as this makes me think of strains; better to say "changes", no?). These changes include snps, insertions, deletions, and "complex"… no idea what complex is? The figure legends are completely insufficient. And I still don't know if you are tallying in some kind of number of recombinations and psuedogenizations into the mix (I assume these are included in the frame-shifts)? The quantification is murky to me.

We used third codon positions mainly to facilitate comparison with other studies; e.g., the Richardson et. al analysis of *Wolbachia* evolutionary rates (https://doi.org/10.1371/journal.pgen.1003129). It is however common to only use mostly neutrally evolving sites to determine evolutionary rates in order to avoid differences arising from adaptive processes.

The figures the reviewer is referring to aim to convey different types of information: Figure 2 displays the evolutionary rate estimates from neutral sites in comparison to other symbionts and pathogens. Figure 3 in contrast displays all changes we observed in a single strain of *Spiroplasma*.

The adhesin proteins are evolving fast. But aren't Spiroplasma commonly intracellular… so why would it be binding an extracellular protein? … can you discuss this? I presume invasion or something?

*Drosophila*-associated *Spiroplasma* are mostly extracellular, although they experience an intracellular phase during vertical transmission when they infect oocytes. We know that in other *Spiroplasma* species, adhesins are involved in insect cell invasion (https://doi.org/10.3389/fcimb.2017.00013, https://doi.org/10.1371/journal.pone.0048606) and we have now clarified this in the discussion:

"For example, adhesion-related proteins are important in cell invasion in other *Spiroplasma* species (Béven et al., 2012; Dubrana et al., 2016; Hou et al., 2017) and are enriched for evolutionary changes in sHy and sMel (Fig. 2)."

There might be a correlation with genome size and speed of evolution. You mention this in the discussion, but briefly. Can you elaborate on this, especially because Spiroplasmas are close to mycoplasmas which are REALLY small genomes.

There is some novel evidence that prokaryotic genome size is strongly correlated with mutational rate (https://doi.org/10.1016/j.cub.2020.07.034), rather than mostly determined by effective population size as previously suggested. This novel study also found that increased mutation rates often occur in lineages that have lost DNA repair genes, which is in line with our findings. Comparing evolutionary rates (Fig. 1) with genome sizes and the presence of DNA repair genes reveals that correlation is not straightforward for the endosymbiotic lineages we compared. For example, *Wolbachia* and *Buchnera* appear to have lower substitution rates than *Spiroplasma*, yet both have ~similar genome sizes (*Wolbachia*) or smaller genomes (*Buchnera*) than *Spiroplasma poulsonii*. We have included the discussion on mutational rates determining genome size as follows:

"Further to absence of DNA repair genes causing elevated mutation rates, a recent comparative study demonstrated a strong negative correlation between mutation rate and genome size in free living and endosymbiotic bacteria (Bourguignon et al., 2020). This correlation is however not apparent in the genomes of endosymbionts we have investigated. For example, the considerably slower evolving *Buchnera* genomes are much smaller than *Spiroplasma*, and *Wolbachia* would be predicted to have much larger genomes if their size was mainly determined by mutational rates. This suggests that mutational rates alone are a poor predictor for the sizes of the here investigated genomes. Likely, these genome sizes result from an interplay of multiple factors such as population size, patterns of DNA repair gene absence, and mutational rates (Kuo et al., 2009; Marais et al., 2020)."

We have further moved supplementary Figure S5 into the main manuscript body (now Fig. 7) to better enable the readers to follow the discussion on the lack of DNA repair genes.

Figure 3 is really confusing. I assume FS is frameshift, is IF induced fragmentation? After about 10 minutes I could decode it. Is this really the best way to think about these results? Perhaps? But perhaps not? ARP? I think it's adhesin stuff, but you don't say this until later.

We have revised and clarified all figure legends. Please also see the comment above.

### Reviewer #2:

General assessment:

This work utilizes two Spiroplasma populations as the materials to study the substitution rates of symbiotic bacteria. A major finding is that these symbionts have rates that are ~2-3 orders higher than other bacteria with similar ecological niches (i.e., insect symbionts), and these substitution rates are comparable to the highest rates reported for bacteria and the lowest rate reported for RNA virus. Based on these findings, the authors discussed how this knowledge could be used to infer and to understand symbiont evolution. The biological materials used (i.e., symbionts maintained in fly hosts for 10 years and cultivated outside of the host for > 2 years) are valuable, the technical aspects are challenging, and the answers obtained are certainly interesting. The key concern is the limited sampling of other bacteria for comparison to derive the conclusions.

Major comments:

1) The key concern regarding sampling involves several points. (a) The two populations represent the species Spiroplasma poulsonii. Is this species a good representative for the genus? Or is it an exception because it is a vertically inherited male-killer? Most of the characterized Spiroplasma species appear to be commensals and are not vertically inherited. (b) The other species with a comparable rate is Mycoplasma gallisepticum (i.e. a chicken pathogen that spreads both horizontally and vertically). Mycoplasma is a polyphyletic genus with three major clades. While closely related to Spiroplasma, their hosts and ecology are quite different. Do all three groups of Mycoplasma have such high rates? If so, are the high rates simply a shared trait of these Mollicutes and has nothing to do with the distinct biology of S. poulsonii? How about other Mollicutes (e.g., Acholeplasma and phytoplasmas). (c) The group "human pathogens" in Fig. 2 show rates spreading across four orders of magnitude. This is too vague. How many species are included in this group? Are their rates linked to their phylogenetic affiliations? (d) Did Fig. 2 provide comprehensive sampling of bacteria? How about DNA viruses? Michael Lynch has done extensive works on mutation rates (e.g., DOI: 10.1038/nrg.2016.104), some of those should be integrated and discussed.

(a) We agree that it is difficult to draw general conclusions of evolutionary rates in the genus *Spiroplasma* from looking at only 2 strains from the same species, and therefore we have not attempted to do so. We also agree that population bottlenecks at vertical transmission events may be a main reason for the elevated substitution rates. In the first version of the manuscript (first section of the discussion), we have therefore focussed our comparisons on Bacteria with similar ecology for which evolutionary rate estimates are available (*Wolbachia*, *Buchnera*, *Blochmannia*).

(b) As far as we are aware, there is some anecdotal evidence that mycoplasmas evolve quickly (https://link.springer.com/article/10.1007/BF02115648) as well as one study estimating evolutionary rates from genome-wide data of multiple *M. gallisepticum* isolates (https://doi.org/10.1371/journal.pgen.1002511). We are unaware of systematic studies estimating evolutionary rates in other mollicutes, and we feel it is beyond the scope of this article to provide such a systematic assessment. However, we do agree that loss of DNA repair genes and elevated substitution rates in *M. gallisepticum* and *S. poulsonii* could also have occurred independently and have now clarified this in the manuscript: “Absence of DNA mismatch repair pathway may thus be ancestral to Entomoplasmatales (Spiroplasmatacea + Entomoplasmataceae) and contribute to the dynamic genome evolution across this taxon (Lo et al., 2016; Rocha and Blanchard, 2002). [New version:] Alternatively, increased substitutional rates caused by the loss of these loci could have arisen multiple times independently in Entomoplasmatales. ”

(c) We have now provided a more comprehensive figure legend that clarifies that the estimate was obtained from 16 different human pathogens. The range provided covers almost the entire mutational spectrum in Bacteria (https://doi.org/10.1099/mgen.0.000094).

(d) Please see the comment under (c). We have now also included an estimate for DNA viruses in Fig. 2.

2) This study is based on two lab-maintained populations. How may the results differ from natural populations? I understand that no estimate may be available for natural populations and additional experiments may not be feasible, but at least a more in-depth discussion should be provided.

We have expanded the discussion on this matter:

"Our rate estimate is potentially biased by at least two factors. First, we have only investigated laboratory populations of *Spiroplasma poulsonii*. Each vertical transmission event creates symbiont population bottlenecks potentially increasing genetic drift and thus substitution rates. Because the number of generations in natural populations of the *Spiroplasma* host *Drosophila hydei* is lower compared with laboratory reared hosts, vertical transmission events are rarer under natural conditions, and substitution rates therefore potentially lower. Further, laboratory strains could experience relaxed selection compared with natural symbiont populations. This may lead to higher substitution rate estimates from laboratory populations compared with natural populations. Secondly, substitution rates often appear larger when estimated over brief time periods (Ho et al., 2005)."

3) The authors use adaptation as a key explanation for several of the findings. Stronger support and alternative explanations are needed. For example, why genome degradation may be used as a proxy for host adaptation (line 497)? If this explanation works only for sHy but not the other strain within the same species (i.e., sNeo), is this still a good explanation? Similarly, for the arguments made in lines 524-528, supporting evidence should be presented in the Results. For example, what are the rate distribution of all genes? Do those putative adaptation genes have statistically higher rates and/or signs of positive selection?

We agree with the reviewer in that we have no direct evidence for adaptation as explanation for the genomic architecture of *s*Hy. We have therefore carefully revised the manuscript to make clear that adaptation is a potential explanation. The key paragraph now reads:

"Using signatures of genomic degradation as a proxy, our findings collectively suggest that *s*Hy is in a more advanced stage of host restriction than *s*Mel. This may indicate host adaptation as a result of the fitness benefits associated with *s*Hy under parasitoid pressure, and the absence of detectable costs for carrying *s*Hy in *Drosophila hydei* (Osaka et al., 2013; Jialei Xie et al., 2014; Xie et al., 2010). However, the *Spiroplasma* symbiont of *Drosophila neotestacea s*Neo is also protective, does not cause obvious fitness costs (Jaenike et al., 2010), but has a less reduced genome (Fig.5, Ballinger and Perlman, 2017). Further, it is also possible that genome reduction in *s*Hy was mainly driven by stochastic effects or even by adaptation to laboratory conditions, as we have not investigated contemporary *s*Hy from wild *D. hydei* populations."

4) The chromosome and plasmids have very different rates (lines 315-316). Since this study aims to compare across different bacteria, perhaps the analysis should be limited to chromosomes for all bacteria.

We have only used chromosomal variants for the rate estimates. From the results section of the first version of the manuscript: "To estimate rates of molecular evolution in *Spiroplasma poulsonii*, we measured chromosome-wide changes in coding sequences of *Spiroplasma* from fly hosts (*s*Hy) and axenic culture (*s*Mel) over time." We now also mention this information in the figure legend for Fig. 2.

5) Formal statistical tests should be performed to test the stated correlations (e.g., lines 360-361, genome size and the number of insertion sequences).

As suggested, we have calculated Pearson's correlation coefficients, which confirm the observation that *Spiroplasma* genome size is correlated with the number of predicted IS elements and proportion of predicted prophage regions (new supplementary file Fig. S4).

6) Fig. 5. The differences in CDS length distribution should be investigated and discussed in more details. The authors stated that they have re-annotated all genomes using the same pipeline, so this finding cannot be attributed to the bioinformatic tools. If these findings are true (rather than annotation artifacts), it is quite interesting. How to explain these? Why is Sm KC3 so different from all others?

There are several potential explanations for the differences in CDS length: 1) The skew towards very short predicted CDS is most pronounced in draft assemblies with relatively many contigs. We therefore think that assembly breaks have resulted in an artificially high number of short CDS by introducing splits mid-CDS. For example, in the Poulsonii clade, the *s*Neo assembly is composed of 181 contigs. This likely explains the higher proportion of very short CDS when compared with *s*Mel and *s*Neo. 2) An excess of short CDS could also indicate many truncated genes that have become pseudogenised. We would therefore expect shorter median CDS lengths in genomes that undergo reduction. In Fig 5, the differences in CDS lengths within the Mirum group may be explained this way: in comparison with *S. eriocheiris*, CDS lengths are shorter for *S. mirum* and *S. atrichopogonis*. The latter 2 genomes also have a lower coding density and genome size, which may indicate recent genomic reduction. 3) Prophage regions are often characterised by shorter CDS, so genomes with overall higher proportions of prophage are expected to harbour a higher amount of smaller CDS. We have added the following statement to the manuscript:

"The distribution of CDS sequence lengths varies across the investigated genomes (Fig. 5), which may be explained by differences in proportion of prophage regions, level of pseudogenization, and assembly quality."

7) Lines 467-479. Multiple lineages have purged the prophages is an interesting hypothesis and may be important in furthering our understanding of these bacteria. More detailed info (e.g., syntenic regions of prophage sites across different species) should be provided in the Results to support the claim. Perhaps the sampling should be expanded to include the Apis clade (i.e., the clade with the highest number of described species within the genus) to test if the prophage invasion occurred even earlier or independently in multiple lineages. Additionally, CRISPR/Cas systems are known to have variable presence across Spiroplasma species (DOI: 10.3389/fmicb.2019.02701). How does this correspond to prophage distribution/abundance?

For *s*Mel, none of the prophage regions predicted with PHASTER show clear synteny over the majority of their length in *s*Hy, which makes synteny comparison (including across even more distantly related strains) difficult. CRISPR-Cas systems are entirely absent in Citri and Poulsonii clades, so are unlikely to be responsible for differences in prophage proportions between *s*Mel and *s*Hy. For the revised version of the manuscript, we have performed two additional analyses focussing on prophages and CRISPR/Cas in *Spiroplasma*, and have expanded the sampling to the Apis clade, as suggested by the reviewer.

Firstly, we have investigated the history of prophage-related loci across the *Spiroplasma* phylogeny. Gene tree - species tree reconciliations suggest that the number of prophage loci have expanded greatly in some of the lineages, especially in the Citri clade. Many of these expansions have happened relatively recently, and therefore most likely occurred independently in multiple lineages.

Secondly, we have used two approaches to predict CRISPR/Cas systems and arrays. We found CRISPR/Cas systems, or their remnants only in the Apis clade, which coincides with the absence of prophage loci in most members of this clade. Based on Cas9 phylogeny, there were multiple origins and several losses of Cas9 systems in the Apis clade. Interestingly, in some taxa with reduced Cas9 systems (e.g., *S. atrichopogonis* and *S. mirum*), there are elevated numbers of phage loci which suggests that phage invasion in *Spiroplasma* is linked to the loss of antiviral systems, as has been suggested previously.

Overall, these data are in line with *Spiroplasma* being susceptible to viral invasion when CRISPR/Cas is absent. Highly streamlined genomes in the absence of

CRISPR/Cas might thus be explained by loss of prophage regions or by a lack of exposure to phages. We have revised the paragraph discussion prophage distribution:

"It was therefore argued that phages have likely invaded *Spiroplasma* only after the split of the Syrphidicola and Citri+Poulsonii clades (Ku et al., 2013). Our prophage gene tree-species tree reconciliations are in line with this hypothesis, but also indicate that prophage proliferation has largely happened independently in different *Spiroplasma* lineages (Fig. S4, supplementary material). CRISPR/Cas systems have multiple origins in *Spiroplasma* (Ipoutcha et al., 2019) and only occur in strains lacking prophages (Fig. S4, supplementary material). While the absence of antiviral systems often coincides with prophage proliferation (e.g., in the Citri clade), several strains with compact, streamlined genomes lack CRISPR/Cas and prophages (e.g., TU-14, Fig. S4, supplementary material). These strains also show other hallmarks of reduced symbiont genomes (small size, high coding density, lack of plasmids and transposons, Fig. 5), which is in line with the model of genome reduction discussed above and suggests prophage regions were purged from these genomes. Alternatively, these strains may never have been exposed to phages."

Minor comments:

1) Lines 32, 517, and possibly other parts: Use "increased" or "decreased" to describe the rate differences are inappropriate because these imply inferences of evolutionary events after divergence from the MRCA, which are clearly not the case. It would be more appropriate to use "higher" or "lower" to describe the difference.

We agree and have revised the use of these terms. In the new version of the manuscript we only use 'increase' or 'decrease' 'when we refer to a change compared with MRCA.

2) Lines 31-32. This is too vague. For the rates, the description should be more explicit (e.g., higher by X orders of magnitude). The term "symbiont" is also vague. Broadly speaking, all human pathogens (included in Fig. 2) or plant-associated bacteria could be considered as symbionts as well. Would be better to define this point more clearly.

Corrected:

"We observed that *S. poulsonii* substitution rates are among the highest reported for any bacteria, and around two orders of magnitude higher compared with other inherited arthropod endosymbionts."

3) Fig. 1. The alignment is off. For example, June should be located near the middle between two tick marks.

The tick marks did not correspond to year boundaries. We recognise that this may be confusing and have adjusted the image for the new version of the manuscript.

4) Line 207. This is confusing. There should not be 6 circular chromosomes.

Corrected 'chromosomes' to 'contigs'.

5) Line 211. Why is the hybrid assembly more fragmented?

The hybrid assembly algorithm used by Unicycler (https://doi.org/10.1371/journal.pcbi.1005595) first creates an assembly from the short reads and then uses long reads to span repeats and other questionable nodes in the assembly graph. We suspect that if the initial short read assembly is highly fragmented (such as is the case for *S. poulsonii*), even a large amount of high quality long reads cannot fully resolve the assembly graph. Our approach was therefore to use the complete long read assembly as starting point.

6) Methods and Results. More detailed information regarding the sequencing and assembly should be provided. For example, how much raw reads were generated for each library? What are the mapping rates? How much variation in observed coverage across the genome?

We now provide these details in the new Supplementary table S2.

7) Lines 341-342. How to establish an expected level of synteny conservation?

We have removed the reference to 'expected' levels of synteny.

8) Line 487. I do not see how this statement could be supported by Fig. 5. Also "less pronounced" is vague.

Corrected to

"However, when using the similarity agnostic tool PhiSpy, the predicted prophage regions were similar in size between *s*Hy and *s*Mel (Fig. S2)."
